# SIDERITE: Unveiling Hidden Siderophore Diversity in the Chemical Space Through Digital Exploration

**DOI:** 10.1101/2023.08.31.555687

**Authors:** Ruolin He, Shaohua Gu, Jiazheng Xu, Xuejian Li, Haoran Chen, Zhengying Shao, Fanhao Wang, Jiqi Shao, Wen-Bing Yin, Long Qian, Zhong Wei, Zhiyuan Li

## Abstract

Siderophores, a highly diverse family of secondary metabolites, play a crucial role in facilitating the acquisition of the essential iron. However, the current discovery of siderophore relies largely on manual approaches. In this work, we introduced SIDERTE, a digitized siderophore information database containing 872 siderophore records with 649 unique structures. Leveraging this digitalized dataset, we gained a systematic overview of siderophores by their clustering patterns in the chemical space. Building upon this, we developed a functional group-based method for predicting new iron-binding molecules. Applying this method to 4,314 natural product molecules from TargetMol’s Natural Product Library for high throughput screening, we experimentally confirmed that 40 out of the 48 molecules predicted as siderophore candidates possessed iron-binding abilities. Expanding our approach to the COCONUT natural product database, we predicted a staggering 3,199 siderophore candidates, showcasing remarkable structure diversity that are largely unexplored. Our study provides a valuable resource for accelerating the discovery of novel iron-binding molecules and advancing our understanding towards siderophores.

## Introduction

Siderophore is a diverse family of secondary metabolites that exhibit high affinities for binding and chelating iron, one of the essential elements for cellular processes including replication and respiration(1,2). However, iron presents a limiting resource due to its low solubility in most environments(3). To overcome this challenge, microbes synthesize small molecule iron chelators, known as siderophores, which are able to bind poorly soluble iron and transport it into the cell via specific transporters(3). The significance of siderophores lies in their vital role in ensuring microbial survival and growth. Pathways associated with siderophore synthesis and uptake are widely present in bacteria, fungi, and archaea(4), constituting complex ecological games(5). Additionally, as a special type of natural product, siderophores exhibit notable antibacterial and antifungal activities, making them promising candidates for the development of novel therapeutics(6). Despite their importance, our current understanding of siderophores is still limited due to their high diversity.

The extensive range of microorganisms lifestyles plays a pivotal role in shaping the remarkable diversity of siderophores. According to a review in 2014, over 500 different types of siderophores have been identified, with 270 having been structurally characterized(7). Despite this wealth of structural information, the number of experimentally characterized siderophore biosynthetic pathways remains considerably lower(3,8–10). Siderophore biosynthesis predominantly falls into two major classes of pathways: the non-ribosomal peptide synthetase (NRPS) pathway and the NRPS-independent siderophore synthetase (NIS) pathway(3,11). Notably, the NRPS pathway has been recognized for its capacity to yield a diverse range of natural products(4). With such a high diversity, a systematic overview of siderophores, encompassing their sources within producing organisms, biosynthetic pathways, chemical properties, and a quantitative assessment of their diversity within the known chemical space of natural products, is still to be developed.

Thanks to the efforts of countless researchers over the past few decades, significant progress has been made in systematically analyzing siderophores. In 2010, Robert C. Hider and Xiaole Kong provided a valuable source of information on the chemistry and biology of siderophores in a seminal review, which included structural features of 294 siderophores in appendix(3). A siderophore database, known as the “siderophore base” (http://bertrandsamuel.free.fr/siderophore_base/index.php), was created by Samuel Bertrand in 2011, which contained 262 siderophores with structure images, organism sources, and references. However, the database was last updated in 2013 and is no longer maintained. Moreover, new siderophore molecules (12–15), and even new siderophore functional group types (diazeniumdiolate and 2-nitrosophenol)(16–19), are constantly being discovered in various microorganisms.(3) Information regarding these siderophores is currently dispersed across various publications and needs to be systematically recorded.

Another significant challenge in achieving a systematic overview of siderophores is the lack of digitalization, which hinders computational investigations. In the field of natural products, large digitized databases such as COCONUT(20), LOTUS(21), and SuperNatural(22) record their molecules in SMILES (Simplified Molecular Input Line Entry System) format. SMILES is the commonly used format for storing and analyzing chemical molecules, which translates a chemical structure into a string of symbols that are easily readable by computer softwares(23). This format enables large-scale computational investigations such as machine learning(24). However, there is no systematically curated digital dataset about siderophores. Digitalized natural product databases do not offer publicly accessible and searchable instances of "siderophore" and only contain a fraction of currently known siderophores. Previous case-by-case research on siderophores reported their siderophore discoveries by images of the structural formula(12–16), as do previous reviews and the siderophore base. Although tools are available for converting chemical structure images to SMILES, their accuracy is limited(25–27). Furthermore, the images in previous reviews and Samuel Bertrand’s siderophore base are not standardized and have some errors, further complicating the conversion(3). Therefore, in addition to curating the comprehensive information set of siderophores, digitalization also constitutes a significant challenge.

With advances in siderophore research and computational biology, the time is opportune for a quest for answers. For instance, how shall we quantify the chemical similarity between siderophores to understand whether the iron scavenging systems are under convergent or divergent evolution? And practically, can we identify new compounds with iron-binding abilities from the vast repository of natural products? Given the crucial role of iron in microbial metabolism(1), identifying such iron-binding compounds would facilitate targeted interventions in microbial communities(6,28) and guide therapeutic applications (iron chelation therapy)(29). Suppose a novel family of iron-chelating siderophores that can be utilized by beneficial but not pathogenic microbes. These chemicals can serve as targeted prebiotics(5,30,31). Furthermore, iron homeostasis plays a critical role in human health and is implicated in various conditions ranging from acute iron poisoning to cancer and aging(32–34). Unfortunately, medicines capable of modulating iron homeostasis remain scarce(35). Therefore, can we draw inspiration from siderophores and investigate the iron-binding potential of the vast array of currently available small molecules? To address these questions and many more, a comprehensive quantification of the currently known siderophores repository is necessary.

Taken together, establishing a comprehensive siderophore database is crucial for gaining a deeper understanding of siderophore synthesis, function, and application. To fulfill this need, we have developed the Siderophore Information Database (SIDERTIE), a user-friendly platform that includes 872 siderophore records with 649 unique structures in the SMILES format, covering all known siderophores up to May 2023. Leveraging SIDERTIEs digitalization capabilities, we presented the most comprehensive statistics of siderophores to date, covering biosynthetic pathways, source of producers, and several chemical characterizations. The dispersed distribution of known siderophores within the chemical landscape of natural products hints at the vast, largely uncharted territory of undiscovered diversity. Building upon this quantitative overview, we proposed a functional group-based method to batch discover new siderophores. A total of 48 siderophore candidates were identified from 4,314 natural product molecules from TargetMol’s Natural Product Library by this method, 40 of which were experimentally confirmed to possess iron-chelating properties. The creation of the siderophore database and the swift discovery of new siderophores hold profound implications for advancing siderophore-related scientific research.

## Result

### 1. Overview of SIDERITE

The Siderophore Information Database (SIDERITE, http://siderite.bdainformatics.org) contains 872 records covering all known siderophores up to May 2023. In addition to siderophore records from previous databases and reviews(3,11,36–38), 224 records were curated from single research articles for the first time (Table S1). In addition to the expanded collection, SIDERITE records the siderophore structures in the SMILES format, a string-based digitization of chemical structures that can be easily processed by various algorithms. The digitized siderophore structures provide a more convenient way of presenting the information compared to previous databases and reviews, which primarily used the picture format. Overall, SIDERITE represents a valuable resource and facilitates systematic computational investigations. Notably, in comparison to other siderophore collections, SIDERITE boasts the largest collection of siderophores and stands out for being freely accessible and digitized (Table 1).

**Table 1.**
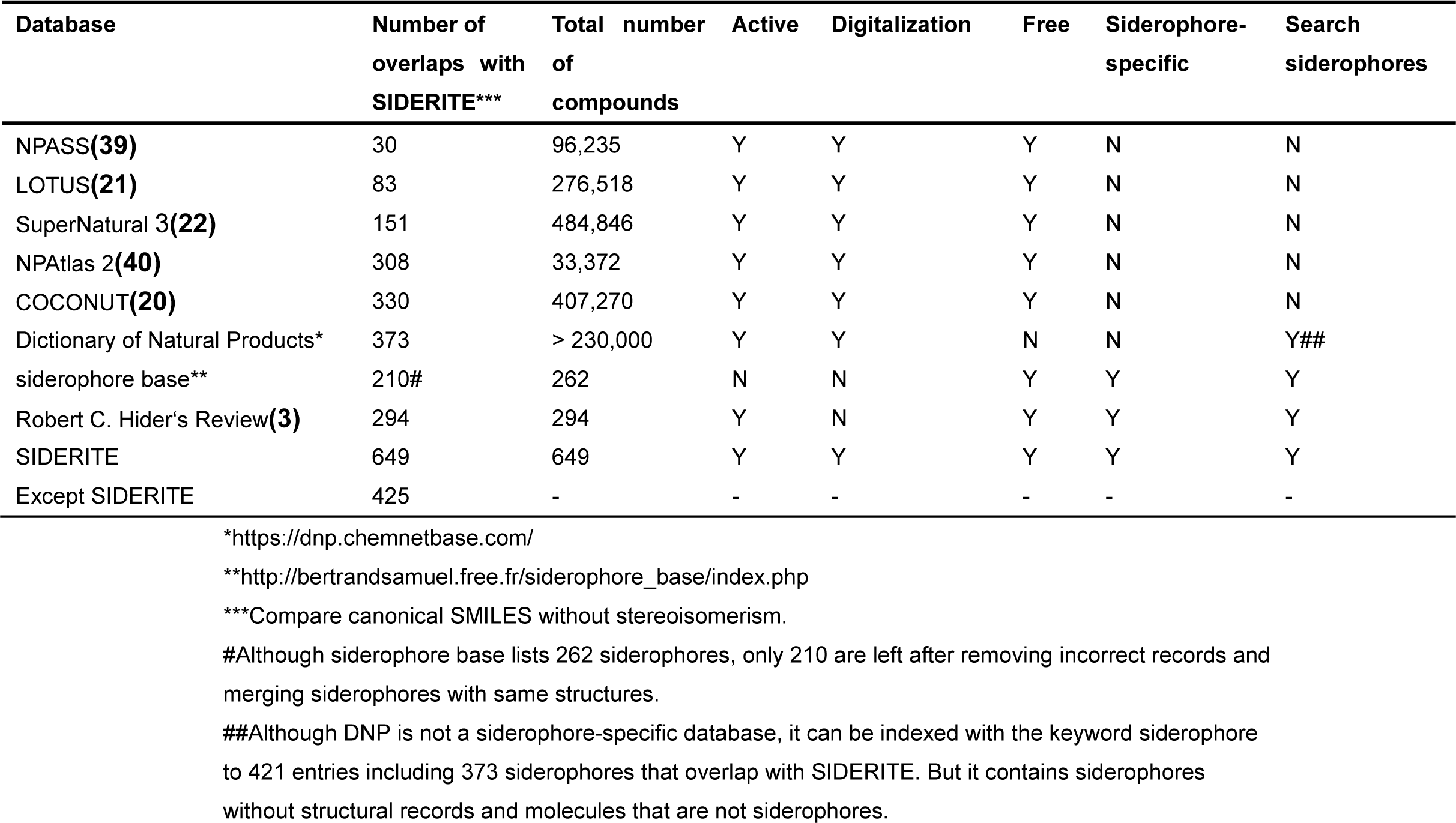
Comparison of databases related with siderophores.

Digitizing siderophores enables computational analysis, particularly in unifying siderophores based on their chemical structures. By comparing the canonical SMILES of siderophores, we identified 649 unique siderophore structures out of the 872 total records (Table S2). During this process, we observed that many siderophores share identical structures but have different names, such as acyl-desferrioxamine 1 and legonoxamine A, bacillibactin and corynebactin, streptobactin and griseobactin, and brucebactin and lysochelin. This observation indicates that the same siderophores were discovered in different species or by different research groups(41–48). Therefore, for each unique siderophore structure, we merged corresponding records and designated one of their names as the official “Siderophore name”, while recording the other names as “Siderophore other name”.

The 649 unique siderophores in our database can be classified by their producer sources (Fig 1A). At the kingdom level, the majority of 649 siderophores in the database are produced by bacteria (85.90%), followed by fungi (12.40%), plants (1.54%), and animals (0.15%). While most siderophores are specific to one producer source at the kingdom level, there is one exception. Triacetylfusarinine can be produced by both bacteria (*Paenibacillus triticisoli*) and fungi (*Penicillium* sp.). At the phyla level, 872 producers of 649 siderophores are spread across five major bacterial phyla (229 in Actinobacteria, 7 in Cyanobacteria, 33 in Firmicutes, 412 in Proteobacteria, and 8 in Bacteroidetes) and three major fungal phyla (124 in Ascomycota, 38 in Basidiomycota and, 8 in Mucoromycota, Table S1).

**Figure 1.**
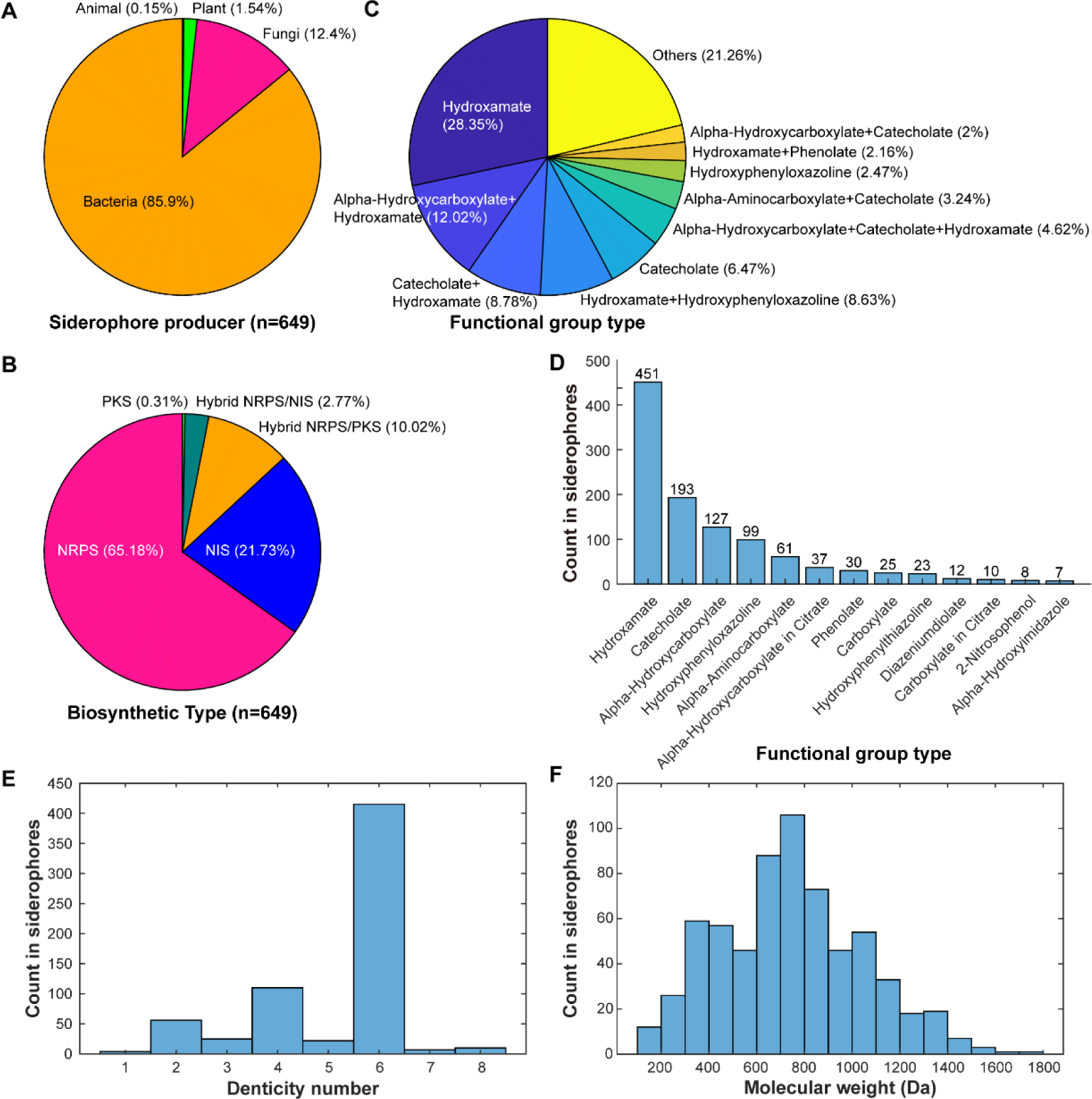
The statistics of 649 unique siderophores in SIDERITE. **A.** Distribution of the siderophore producers by their kingdoms. **B.** Distribution of the siderophore biosynthetic pathways. **C.** Distribution of the functional group type combinations. For clarity, only the top ten combinations are shown, and the others are merged into "Others". **D.** Distribution of the common functional group of siderophores. One siderophore could contribute to more than one functional group type if it contains many types of functional groups. **E.** Distribution of denticity numbers. **F.** Distribution of the molecular weight.

Siderophores can also be classified by their biosynthetic pathways (Fig 1B). Previous studies have suspected that the NPRS pathway is more dominant for siderophore synthesis(3), yet no statistically concrete ratio between NRPS and NIS-derived siderophore has been established. In SIDERITE, we found that related pathways can be classified into NRPS (65.18%), NIS (21.73%), Hybrid NRPS/PKS (10.02%), Hybrid NRPS/NIS (2.77%), and PKS (0.31%). Consistent with previous studies, NRPS was indeed the most abundant biosynthetic pathway of siderophores, followed by NIS. PKS siderophores are rare, with only two cases (proferrorosamine A and tetracycline). In different kingdoms, the composition of the biosynthetic pathway differs (Table S2). In plants and animals, all siderophores are synthesized by NIS. In fungi, almost all siderophores are synthesized by NRPS (90.12%, 73/81). In bacteria, the diversity of the siderophore biosynthetic pathways is higher, with 62.90% (351/558) NRPS, 22.58% NIS (126/558), 11.29% hybrid NRPS/PKS (63/558), and 2.87% hybrid NRPS/NIS (16/558).

Digitization also enables us to easily access the statistical properties of siderophores, such as functional group distribution. Siderophores chelate iron by several common functional groups (coordinating groups)(3), and a single siderophore may use multiple types of functional groups. In SIDERITE, the top five combinations of functional groups are present in 64.25% of the siderophores (Fig 1C). These top five combinations are Hydroxamate (28.35%), Alpha-Hydroxycarboxylate+Hydroxamate (12.02%), Catecholate+Hydroxamate (8.78%), Hydroxamate+Hydroxyphenyloxazoline (8.63%) and Catecholate (6.47%). Hydroxamate and Catecholate are the most common functional groups found in 69.49% and 29.74% of siderophores, respectively (Fig 1D). Most of siderophores have three bidentate groups which forming octahedral geometry with iron in coordinating number six(3). As expected, most siderophores have a denticity number of six (Fig 1E). However, there are exceptions, such as pacifibactin, desferrioxamine T1, malleobactin D and pyoverdine 7.7, which contain four bidentate groups. It’s reported that pacifibactin coordinates iron in a 1:1 ratio, and the function of the extra coordinating group is unknown(49).

The molecular weight (MW) is another important property of siderophores, as it influences the diffusion of siderophores(50). Siderophores are typically small molecules, and the MW of siderophores in our database ranges from 138.12 Da (salicylic acid) to 1766.86 Da (pyoverdine IB3). Most siderophores (90.60%, 588/649) have a middle MW ranging from 300∼1100 Da (Fig 1F). Of note, half of the smallest 12 siderophores (MW < 200 Da) are monomers of other siderophores, such as salicylic acid, 2,3-dihydroxybenzoic acid, and citrate. Most of the large siderophores (MW > 1200 Da) are pyoverdines (87.76%, 43/49), as pyoverdines are frequently composed of more than ten amino acids. Further, we check whether MW distribution is related to biosynthetic types. We found NRPS siderophores exhibit a wide range of MW, from 206.20 Da (Spoxazomicin D) to 1766.86 Da (pyoverdine IB3). NRPS siderophores are generally heavier (mean: 835.68 Da; median: 816.00 Da) than NIS siderophores (mean: 506.93 Da; median: 516.63 Da).

Additionally, aqueous solubility and the diffusion coefficient also are important ecological properties of siderophores. However, most siderophores have not been measured experimentally for these two properties. Therefore, we employed SolTranNet(51) and Stokes–Einstein Gierer-Wirtz Estimation (SEGWE)(52) to predict these properties using SMILES representations (Fig S1). Out of the total 649 siderophores, 635 were predicted to be water-soluble (predicted logS>-6). Among them, 551 siderophores exhibited a “good” level of aqueous solubility (predicted logS>-4). Notably, 14 siderophores exhibit poor aqueous solubility (predicted logS≤-6) due to the presence of long-chain fatty acids, suggesting them are probably insoluble in water. The minimum and maximum predicted diffusion coefficients are 2.66×10^-10^ m^2^/s and 7.7×10^-10^ m^2^/s respectively. The majority of siderophores (611 out of 649) exhibit predicted diffusion coefficients within the range of 2.66 to 5.50×10^-10^ m^2^/s, consistent with previous research(53).

### 2. Clustering of siderophores by their structural similarities

Siderophores have been known to exhibit remarkable structural diversity(3) (Fig S2). Converting siderophores into SMILES format enables us to quantify their chemical similarity more effectively, both within the SIDERITE database and between other natural products. To systematically assess the structural diversity of siderophores, we first locate all 649 SIDERITE structures to the vast chemical space of the COCONUT database by merging all molecules in these two databases, which encompasses over 4×10^5^ natural products. By Tree MAP (TMAP) visualization of chemical similarity (Fig S3), we observed that the 649 siderophores could be grouped into 25 distinct clusters, which were separated from each other by natural products from the COCONUT. The clustering result shows siderophores have unevenly distributed structural diversity (Fig 2, Table S2). Most of these clusters (16 out of 25) only contain a few members (<5), while the largest four clusters account for 89.37% of the siderophore structures in total. We sorted the cluster indexes by their member counts in descending order; for instance, cluster 1 contains the most siderophore structures.

**Figure 2.**
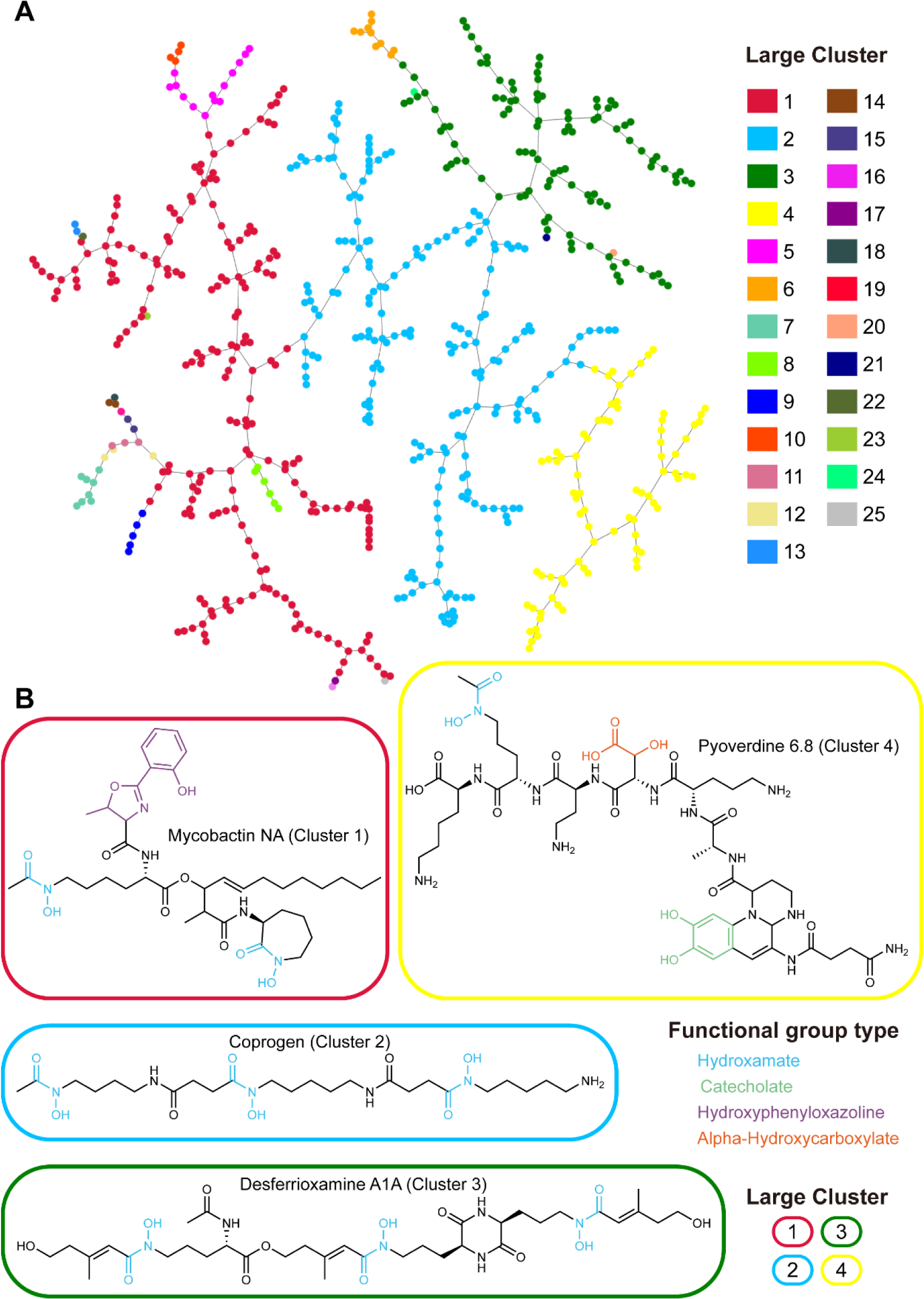
The visualization of large siderophore clusters in SIDERITE. **A.** A network of siderophores connected by chemical similarities. Each node in the network corresponds to a siderophore molecule, and the nodes are linked to their most similar neighbors, forming a minimum spanning tree(57) (described in the method section). Nodes are colored by their cluster IDs. **B.** For the four largest clusters, example structures are provided, and the functional groups of the siderophores are colored according to their types. Siderophores are circled by rounded rectangles to show their cluster IDs (same color codes as (**A**)).

Within each cluster, there are common features of functional groups or biosynthetic types. Cluster 1 (201, 30.97%) includes siderophores with phenyl ring structures in the functional groups such as catecholate, phenolate, hydroxyphenyloxazoline, and hydroxyphenylthiazoline. Siderophores in the cluster 1 are synthesized by both NRPS and NIS. Cluster 2 (197, 30.35%) only includes siderophores produced by NRPS pathways except Albomycins by the hybrid NRPS/NIS pathway. Albomycins are naturally occurring sideromycins (siderophore–antibiotic conjugates) produced by some streptomycetes, and their siderophore parts are synthesized by NRPS. From the perspective of functional groups, most siderophores in the cluster 2 contain hydroxamate of (92.39%, 182/197), and many also contain alpha-hydroxycarboxylate (37.06%, 73/197). Cluster 3 (103, 15.87%) is all NIS siderophores. Like cluster 1, most of them contain hydroxamate (90.29%, 93/103), and many also contain alpha-hydroxycarboxylate (33.01%, 34/103). The sources of alpha-hydroxycarboxylate are mostly citrate. Cluster 4 (79, 12.17%) is NRPS siderophores with chromophores such as pyoverdines (93.67%, 74/79). Other small clusters all are located on the edge of four large clusters (Fig 2A). They consist of similar functional group composition (Fig S4-S16), which indicates the possibility of unusual siderophores evolving from common siderophores.

Molecule weight (MW), carbon-nitrogen ratio (C/N), and carbon-oxygen ratio (C/O) of siderophores pertain to their biosynthetic cost in various aspects: higher MW can indicate more building blocks; while producing products with lower C/N or C/O could influence growth under nitrogen-limited or hypoxic environments(54–56). Therefore, we investigated these properties of siderophores in different clusters (Fig S17). Siderophores in cluster 4 have significantly higher MW than average, as most cluster members are pyoverdines. Regarding C/N and C/O ratios, different clusters also exhibit different properties. For example, siderophores in cluster 7 and cluster 10 have significantly higher-than-average C/N because they only contain 1∼2 nitrogen atoms, and siderophores in cluster 4 have a significantly lower-than-average C/N (Fig S18).

Large clusters only provide an initial classification relative to other natural product molecules. Further, we defined small groups within each of the 25 clusters by their structure similarity coefficient (Dice coefficient, threshold 0.6, see method), and obtained 102 small groups in total. Each group is named by its cluster ID *x* and its group ID *y* within this cluster. Accordingly, each siderophore is assigned a unique id *x.y.z*, where z stands for the z-th record within the group. For example, enterobactin has the id 1.22.2. In this ID, “1” means the 1^st^ large cluster. “22” means the 22^nd^ group within the 1^st^ large cluster. “2” means the 2^nd^ record in the 1.22 cluster. In the future, newly discovered siderophores will also be assigned a unique id when being incorporated into the SIDERITE database.

This cluster-group naming method provides an intuitive framework for comprehending the siderophore clusters in the chemical space. The distribution of structural diversity of siderophores in the chemical space is uneven. The three largest clusters contribute to 70.59% (72/102) of the groups, while the average member number of the remaining groups is 1.36 (Table S2). All small groups encompass siderophores derived from unique biosynthetic types, except groups 1.5 and 1.10, where siderophores are synthesized through both hybrid NRPS/PKS and NRPS pathways. The siderophores in these two groups have two biosynthetic types due to the optional modification from PKS. Also, siderophores within the same group are typically produced by species from the same kingdom. However, groups 2.1, 2.9, 3.7, and 9.1 contain producers from both bacteria and fungi, and group (11.1) is produced by bacteria and animals.

For siderophores with unknown biosynthetic types, their biosynthetic types could be inferred by other members with known biosynthetic types in the same small cluster because almost all members in the same small cluster have the same biosynthetic types (except for 1.5 and 1.10 clusters). It’s useful for mining their biosynthetic genes in the genome after discovering new siderophores, which would accelerate siderophore research from structures to genes.

### 3. Discovering new siderophores by the functional group-based method

The clustering analysis of known siderophores has unveiled the presence of common functional groups among these compounds. Drawing inspiration from this observation, we have introduced an innovative rule-based approach, aimed at the discovery of novel siderophores by their chemical structures (Figure 3A-3C). In this approach, we first distilled 15 common functional groups derived from the characteristics of known siderophores. Any molecule containing at least one of these 15 functional groups is identified as a potential siderophore (Figure 3D and Table S3). Then, we exclude candidates containing any of the 8 modified siderophore functional groups incapable of forming coordination bonds to chelate iron (Figure 3E and Table S4). Besides, while the alpha-hydroxycarboxylate functional group is prevalent within siderophores (Figure 3F), we have excluded it from our rule set due to its ubiquity in non-siderophore molecules.

**Figure 3.**
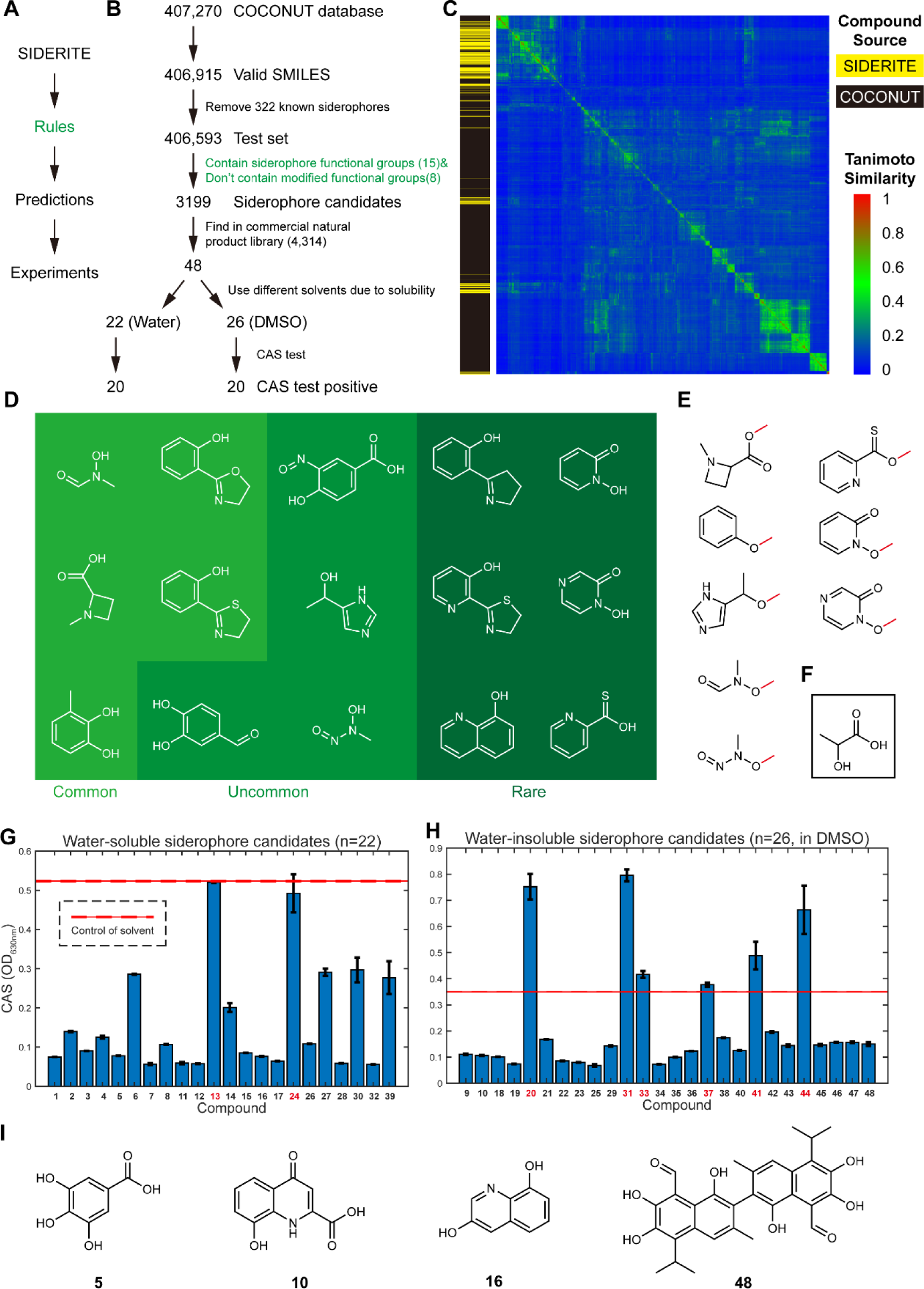
The rule-based siderophore discovery approach and the result of CAS test experiments. **A.** Principle of rule-based siderophore discovery approach. The rules are summarized from known siderophores based on the SIDERITE database. The predictions were then tested by experiments. **B.** Pipeline of the functional group-based siderophore discovery. Molecules containing at least one of 15 functional groups in (**D**) and not any of the 8 modified functional groups in (**E**) are selected as siderophore candidates. **C.** The structural diversity of new potential siderophores in the COCONUT database. 3,199 molecules with potential iron-binding activities and 649 known siderophores were clustered with Tanimoto similarity. The source of molecules (COCONUT or SIDERITE) is shown in the left bar by yellow and black colors, respectively. **D.** The structures of siderophore functional groups in the rules in **(B)**. The rarity of functional groups is noted by the different background colors. **E.** The structures of modified functional groups in the rules in **(B)**. Modifications that cause functional groups to lose iron-chelating abilities are marked in red. **F.** The structure of common functional group alpha-hydroxycarboxylate. **G.** The CAS result of the 25 water-soluble siderophore candidates. The solvent is water. The mean and ± one standard deviation of OD (630nm) are shown in the bar graph. The OD (630nm) of solvent is marked as a red line (mean value) and red dash line (one standard deviation). The compounds with negative results in the CAS test are marked in red. **H.** The CAS result of 30 water-insoluble siderophore candidates. The solvent is DMSO. Other legends are the same as (**F**). **I.** The representative examples of compounds with iron-binding activities.

To verify this method, we applied our functional group-based method to the large chemical database COCONUT, excluding 322 known siderophores from consideration. Remarkably, this analysis led to the identification of 3,199 molecules exhibiting potential iron-binding activities from a database containing over 0.4 million compounds (Table 1 and Table S5). By the Tanimoto similarity clustering (Figure 3C), a large proportion of these potentially iron-binding molecules is scattered in the chemical space, not close to any of the known siderophores. Specifically, only 284 out of the 3,199 molecules exhibit significant structural similarity (with a maximum Tanimoto similarity of > 60%) to the known 649 siderophores cataloged in SIDERITE. The remaining 2915 molecules are strong candidates for novel siderophores with relatively unexplored chemical structures. This analysis underscores the notion that the structural diversity of siderophores remains largely concealed, inviting further in-depth exploration and investigation.

Subsequently, we searched for purchasable molecules out of the 3,199 candidates for experimental verification. 48 molecules (Table S6 and Figure 3B) are available in the commercial natural product library (the Natural Product Library for high throughput screening, catalog number L6000, TargetMol, Shanghai, China, June 2023). We checked the producer sources of these 48 molecules and found that, except for chlorquinaldol (compound **29**) which is synthetic(58), the other 47 are natural products derived from plant, insects, microbes, algae, or humans (Table S6). Among these molecules, 22 are soluble in water, while the remaining 26 have poor solubility in water (Table S6). To address this, we dissolved the poorly soluble molecules by dimethyl sulfoxide (DMSO) instead of water. Subsequently, solutions of these 48 molecules were tested by chrome azurol S (CAS) assay, a universal colorimetric method that detects iron-binding molecules(59).

The high positive rate from the CAS assay supports the effectiveness of our functional group-based method. Among the tested molecules, 20 out of 22 (90.9%) water-soluble compounds and 20 out of 26 (76.9%) water-insoluble compounds exhibited iron-binding activity, as evidenced by a noticeable change in the color of the CAS dye (Figure 3G-3H and Table S7). Actually, most molecules with negative CAS results (compound **20**, **24**, **31**, **33**, **37**, **41,** and **44**) exhibited unusual color patterns, which hinders accurate assessment. For instance, their original solutions were significantly dark in color, or reacted with CAS reagents and formed precipitates or turbidity. Actually, compound **24**, **37**, **41**, and **44** did induce color change in the CAS assay solution, but their precipitation interfered with the optical density measurement. Taken together, only one (compound **13**) out of the 48 molecules was confirmed to lack iron-chelating ability.

Of these 40 molecules, which we confirmed to exhibit iron-chelating capabilities (referred to as potential siderophores henceforth), the majority remains largely unexplored regarding their iron-binding potential. Remarkably, only three (compounds **5**, **10**, and **48**, see Figure 3I with their seven derivatives have been previously associated with iron-binding related research, but have not yet been regarded as siderophores: 1. Gallic acid (compound **5**) has been systematically studied for complexation with iron(60) and has been commonly used in the pharmaceutical industry. In these 40 molecules, there are six gallic acid derivates (compounds **12**, **14**, **17**, **23**, **25**, and **27**) with iron-binding activity. 2. Xanthurenic acid (compound **10**) is the precursor of *Pseudomonas* siderophore quinolobactin(61). 3. Gossypol (compound **48**) has been reported as dietary supplementation with interactions with ferrous sulfate(62), implying iron-binding activity. Gossypol derivate gossypol acetic acid (compound **47**) also shows iron-binding activity in our assay. Taken collectively, at least 30 of the molecules we identified may represent novel siderophores.

We then checked the chemical properties of these 40 potential siderophores. Out of the 40 molecules, the most frequent functional group type is catecholate (36/40). Among these, the majority (19/36) are trihydroxy, such as gallic acid (compound 5, Figure 3I). The remaining four molecules not possessing catecholate contain 8-hydroxyquinoline, which is an uncommon functional group. It is worth noting that while it is known that 8-hydroxyquinoline can chelate metal, only two siderophores (quinolobactin and thioquinolobactin) containing this functional group have been reported so far(63,64). Therefore, our discovery of the four molecules significantly expands the list of 8-hydroxyquinoline siderophores. The four molecules are robustine (compound **9**), xanthurenic acid (compound **10**, Figure 3I), jineol (compound **16**, Figure 3I), and chlorquinaldol (compound **29**). Chlorquinaldol is a synthetic compound. Robustine is produced by plant *Zanthoxylum integrifoliolum*(*65*) and *Zanthoxylum avicennae*(*66*). Jineol is produced by the insect *Scolopendra subspinipes*(*67*). The biosynthetic pathways of robustine and jineol are currently unknown.

Regarding the functions of the 40 potential siderophores, most of them (excluding 14 molecules with unknown functions) are known to have antioxidant activities (14/40). This percentage is significantly higher than the proportion of molecules with “antioxidant” annotations in the commercial library (5.1%). The second most common function are anticancer (7/40), followed by anti-inflammatory (4/40, Table S6). The high occurrence of these pharmaceutical functions in potential siderophores motivates us to explore of the large natural product repository for additional molecules with iron-binding capabilities.

### 4. The usage of SIDERITE database

SIDERITE is freely accessible at http://siderite.bdainformatics.org. The current version serves as a platform for searching and displaying known siderophore chemical structures. Users can access desired siderophores by the following three approaches (Fig 4A).

**Figure 4.**
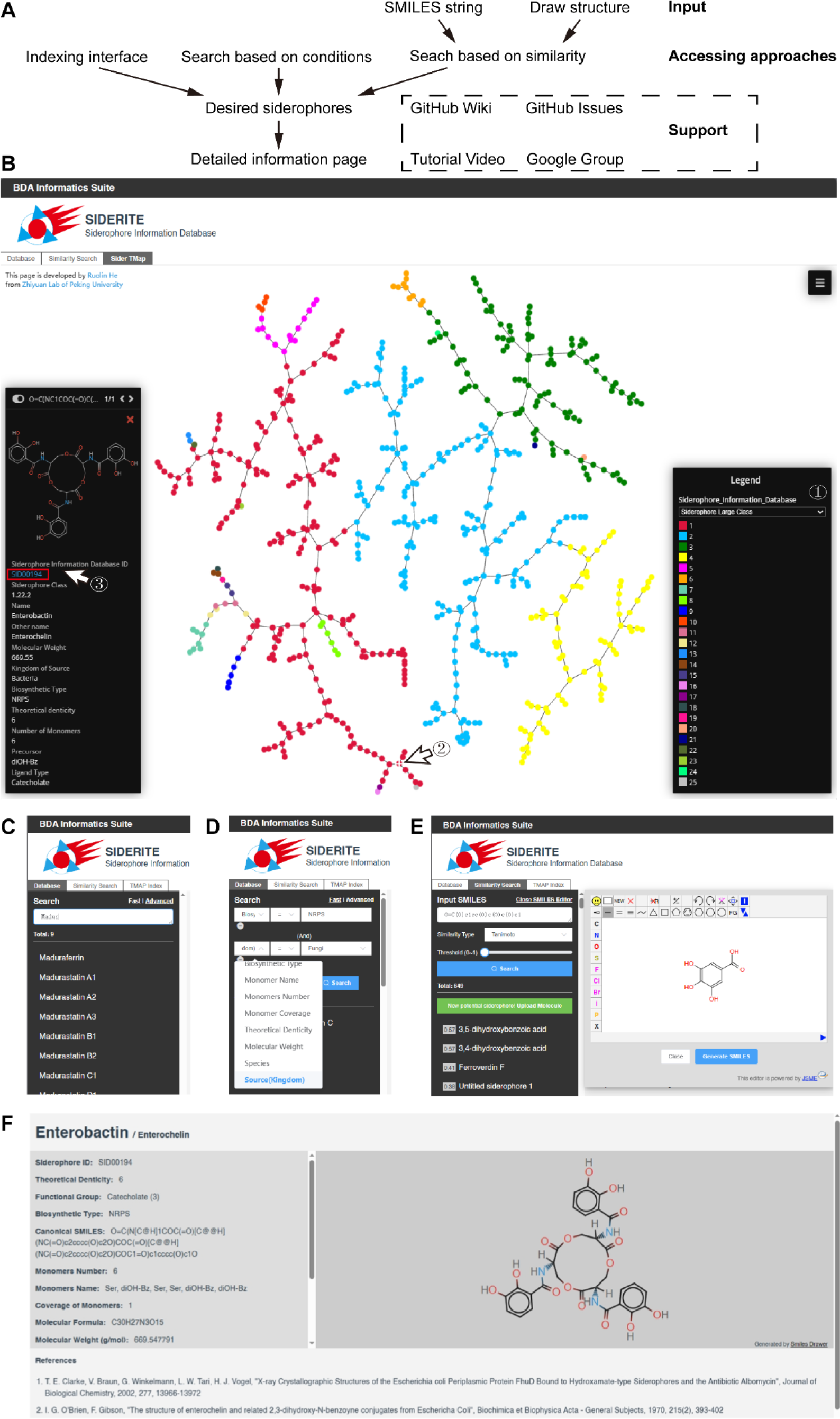
The SIDERITE database. **A.** Workflow of SIDERITE database usage. **B.** Accessing siderophores by the indexing interface by the following steps: 1. Choose a type of siderophore legend. 2. Click the interested siderophore. 3. Jump to the siderophore page by the interactive siderophore ID in the information card. **C.** Fast fuzzy siderophore search. **D.** Advanced siderophore search by precise conditions. **E.** Chemical similarity search, where users can type the SMILES string or draw in the SMILES editor. **F.** The individual page with detailed information on each siderophore structure.

An indexing interface is provided so that users can quickly access the desired siderophores. We visualize the SIDERITE database by TMAP(57), and label multiple properties of siderophores (Figure 2A and Figure 4B, http://siderite.bdainformatics.org/sidertmap). In the mapping, each siderophore is represented as a distinct point that offers more detailed information when clicked. Additionally, users can jump to the siderophore page by the interactive siderophore ID in the information card (Figure 4B).

Users also can use the search function to access desired siderophores (Figure 4C-4D). The platform offers fast fuzzy search functionality based on partial siderophore names, as well as precise advanced search by multiple conditions. The advanced search feature includes various conditions, including siderophore ID, siderophore name, siderophore functional group, biosynthetic type, monomer name, number of monomers, monomer coverage, theoretical denticity, molecular weight, microorganism, source (kingdom), source (phylum), and class. Numeric conditions, such as the number of monomers and molecular weight, can also be set within specified ranges.

Chemical similarity search is the third way to access the siderophore resource in SIDERITE (Figure 4E). For any query molecule, this function helps users to find structurally similar siderophores, and also predict whether this query molecule has iron-binding activity. Users can input the SMILES notation of the query molecule or draw it in the SMILES editor. If the molecule has been predicted to have potential iron-binding activity by our functional group-based method (Figure 3) and is not already present in SIDERITE, the SIDERITE platform will automatically indicate to the user that this is a potentially new siderophore. Users can choose whether or not to upload this molecule to the SIDERITE database without any obligations. Furthermore, this search will return siderophores in SIDERITE that are chemically similar to the query molecule. The chemical similarity metric supports Tanimoto, Dice, and Tversky, and the user can customize the cut-off threshold.

We provide individual pages with detailed information for each siderophore structure (Fig 4F), which provide canonical SMILES, pictures, and related references. Alternative names will be noted after the main name, separated by a forward slash. In the “Functional Group” section, the number of specific functional groups is specified within parentheses. We also provide tutorial materials and feedback channels, which can be accessed in the database or the GitHub Wiki page (https://github.com/RuolinHe/SIDERITE/wiki/Tutorial). Additionally, users can report bugs or pose general inquiries about SIDERITE through GitHub Issues page (https://github.com/RuolinHe/SIDERITE/issues) or our dedicated Google group (https://groups.google.com/g/siderite-database). We are committed to maintaining the SIDERITE database continually and updating it based on the feedback received from our users.

## Discussion

In this work, we have constructed the most comprehensive siderophore database to date (http://siderite.bdainformatics.org/), with digitalized formats. Over the past few decades, a large number of siderophore structures have been identified. Some of them have been collected in Robert C. Hider and Xiaole Kong’s review in 2010(3) and Samuel Bertrand’s unmaintained siderophore database in 2011 (http://bertrandsamuel.free.fr/siderophore_base/index.php). As the field of siderophore research continues to advance, there is a growing need to establish a systematic and standardized classification system for siderophores. While previous dissemination of siderophore knowledge through pictorial formats has undoubtedly facilitated its spread and communication, these formats have not adequately supported systematic computational studies. Based on our digitalized dataset, we were able to obtain a systematic overview of siderophores, and derived an effective functional group-based method for discovering new iron-binding molecules. Our approaches have paved the way for more data-driven discoveries in the field.

Our work provided a statistical overview on siderophores, from the perspectives of biosynthetic type, producer source, and chemical properties. Particularly, the clustering patterns of siderophores in the chemical space hold implications for their eco-evolutionary histories(5), and the drives of their chemical diversity. Siderophores are considered public goods as they are predominantly secretive(68). However, some species have evolved the ability to exploit siderophores without producing them, by synthesizing only the receptors for these siderophores(5,68). To avoid this exploitation, producers can modify the structure of their siderophores(5,69,70). This evolutionary pressure under such a Red Queen situation may generate large siderophore clusters with similar but diverse structures. For example, researchers recently found *Streptomyces venezuelae* produced the uncommon, alternative siderophore foroxymithine (cluster 2.7) to impose a more profound fitness when their main siderophore desferrioxamine (cluster 3.3) was exploited by the yeast *Saccharomyces cerevisiae* (cheater) in the co-culture(71). Meanwhile, siderophores in clusters with few similar members may indicate other strategies, like utilizing multiple structurally distinctive siderophores to avoid intense competition. Across different kingdoms, bacteria exhibit the most diverse synthetic types and the highest number of siderophores, reflecting the intense iron competition observed among prokaryotes(5,49).

Dealing with such a high diversity, one advantages of digitalization is to facilitate the annotation of previously unnoticed siderophores. During our construction of SIDERITE, 50 siderophore records were found by exploring chemically similar natural product molecules around known siderophores on the COCONUT database mapping. Among them, 29 records are from references that do not contain "siderophore" in the title, making their detection through literature searches more challenging. In fact, half of them were initially reported as antibiotics or microbial inhibitors. Also, categorizing siderophores by the SMILES format avoid rediscovering of the same structures from different organisms yet annotating them as “new” siderophores(41–48).

Another advantage of digitalization is to provide resources for data-driven explorations, such as in bio-retrosynthesis prediction. In recent years, deep learning approaches have been employed for predicting the biosynthetic pathways of natural products(72). In addition, high-quality labeled siderophore data can be used to train supervised learning models, leading to the design of novel artificial siderophores with enhanced biological activities.

In fact, our functional group-based method for searching iron-chelating structures serves as a preliminary prototype for data-driven discoveries. Although simple in nature, it can be seen as analogous to supervised learning, where features related to iron-binding (such as rules for including or excluding certain functional groups) are extracted from the existing SIDERITE dataset and applied to chemicals with unknown iron-chelating abilities. With the high successful rate validated by our CAS experiments, this functional group-based method is likely applicable to the vast repository of natural products. In the COCONUT database, our method predicted 3,199 molecules as iron-binding compounds. Particularly, siderophore clusters only containing few members can suggest underexplored siderophore types and highlights for future research. For example, pyridoxatin has only one member in the cluster 18 of SIDERITE. Cordypyridone A and B with antimalarial activity are the nearest natural products to pyridoxatin in the chemical space of COCONUT, and both compounds were predicted as “iron-binding” by our method. However, cordypyridone A and B were not previously considered to be siderophores(^73^), indicating underexplored siderophore types that call for verification by subsequent experiments.

More inquiringly, most of the 3,199 structures are different from known siderophores, suggesting that the true extent of siderophore diversity remains largely unexplored. Our findings also imply a higher prevalence of iron-binding molecules in plants than anticipated. These plant siderophores, known as phytosiderophores, are predominantly produced by graminaceous plants(3). Remarkably, our data demonstrates that 92.5% (37 out of 40) of our identified iron-binding molecules originate from plants (Table S6), revealing substantial untapped potential for phytosiderophores.

Developing new methods to quickly discover new siderophores by systematically summarizing the functional group diversity and structural features of siderophores holds immense scientific and practical importance. Primarily, uncovering novel siderophores can open avenues for innovative therapeutic strategies. Irons critical role in various diseases, including cancer and inflammation, underscores its significance(74–76). It is known siderophores have an antioxidant role by chelating iron because iron can cause reactive oxygen species by reacting with H_2_O_2(77-79)_. And anticancer and anti-inflammatory activities of siderophores are achieved by sequestering iron in the environment to limit the growth of cancer cells(74) and microbes(75,76). More than half of our newly identified iron-chelating molecules have reported antioxidant, anticancer, and anti-inflammatory properties. These molecules may also be used as environmental remediation of heavy metal pollution(80,81).

Different from previous siderophore resources, which remain limited to single studies and therefore do not provide a mechanism to perform cross-study and systematic analyses, SIDERITE provides a curated community-wide platform. All siderophores are digitized, which makes comparisons easy and thus avoids duplicate naming of the same molecule. Each siderophore is assigned a three-level serial unique id based on the siderophore structure similarity feature. The unique id assigned to each siderophore in this study also provides a standardized nomenclature for future studies. With the advent of SIDERITE, newly discovered siderophores can be easily identified and quickly known by other researchers in this field. We invite all relevant researchers to join this community and collaborate to promote the prosperity of the siderophore community.

## Method

### Siderophore information resource

A total of 872 siderophore information records were collected from various sources, including databases, reviews, and research articles. Among these, 355 records were obtained from Samuel Bertrands Siderophore Base, while the remaining records were sourced from other databases, reviews, and research articles (Table S1). Specifically, we obtained 160 new records from the appendix of Robert C. Hider and Xiaole Kongs review, 37 new records from other reviews, 95 new records from the Dictionary of Natural Products (DNP) database (https://dnp.chemnetbase.com/), and 224 new records from research articles by searching with keywords such as "new siderophore," "novel siderophore," and "siderophore discovery". It is important to note that one new record which is a pyoverdine (a type of siderophore) was obtained from the LOTUS database with the keywords “pyoverdine”. This was because this database only allowed searching siderophores by their specific names such as “pyoverdine”, rather than by the general biological function such as “siderophore”.

Each siderophore information record in our dataset comprises various details, including the siderophore name, the synonym name (if any), the type and number of functional groups, the producing species, the biosynthetic type, and the structure (Table S1).

### Biosynthetic type annotation

Information on 721 siderophore biosynthetic types is available in our database, which we documented based on references. For the remaining 151 siderophores with unknown biosynthetic types, we inferred their biosynthetic types by features of different types. The biosynthetic type of the siderophore would be annotated as “Putative NIS”, if it contains monomers that only exist in the NIS siderophore such as citrate, diamine (e.g. 1,3-Diaminopropane, putrescine and cadaverine) or diamine derivatives (e.g. N-hydroxy cadaverine). The biosynthetic type of the siderophore would be annotated as “Putative NRPS” if the monomers in this siderophore all are amino acids.

### Monomer annotation

Monomer annotation of siderophores was applied by rBAN with custom monomer database based on original Norine database(82). Monomers of siderophores are displayed in the individual page of siderophore.

### SMILES conversion from siderophores

The Simplified Molecular-Input Line-Entry System (SMILES) is a notation to describe the chemical structure of molecules using a string format which is particularly useful for subsequent processing by computer. However, in our siderophore information resource, the SMILES format of siderophore structures and their annotations were only available in the DNP database. In contrast, the literature and Siderophore Base only included siderophore structures in the picture format which is more intuitive for readers but poses difficulties for computational analysis. To address this limitation, we developed a customized Python script that utilized the ChemSpider API and Chemical Identifier Resolver (CIR) to convert siderophore names into the SMILES format. The SMILES format structures of 34.93% (124/355) siderophore records in the Siderophore Base were retrieved. For the remaining 748 siderophores that were not found or from other resources, we manually drew their structures and obtained the SMILES format structures by SMILES generator/checker tool (http://www.cheminfo.org/flavor/malaria/Utilities/SMILES_generator checker/index.htm l).

To ensure consistency and facilitate comparison, we then converted all SMILES strings into canonical SMILES strings using another custom Python script. The RDKit package was utilized for this conversion process, while simultaneously obtaining the molecular formula and molecular weight of the siderophores.

### Prediction of aqueous solubility and the diffusion coefficient for siderophores based on SMILES

To predict aqueous solubility, we used a machine learning tool SolTranNet with SMILES of siderophores as input(51). For the prediction of the diffusion coefficient, we used the SEGWE calculator developed by Evans, R. *et al*.(52) with a temperature 298.15K and water as solvent.

### Visualization of siderophore distribution in the chemical space

To compare and analyze siderophores intuitively, we visualize SIDERITE with other chemical databases by TMAP(57). The visualization mapping of SIDERITE with other chemical databases consists of the vast chemical spaces for exploring new siderophores. The TMAP mapping algorithm tends to produce a significant distance between a molecule with and without specification of stereoisomersnon-stereoisomeric counterpart. To avoid this bias, we removed stereoisomerism information based on the canonical SMILES. The intersection of SIDERITE with other chemical databases is removed from the chemical database in the visualization.

### Clustering of siderophores

The identification of large clusters of siderophores was based on their arrangement within the vast chemical space. The chemical space is composed of numerous molecules, and in our study, we employed the COCONUT database(20) which comprises approximately 40.7w small molecules. Within this space, siderophores were found to be grouped into distinct large clusters, separated from other non-siderophore molecules.

To identify smaller clusters within each large siderophore cluster, we utilized a molecular structure similarity threshold of 60%. Specifically, siderophores were considered to be part of the same small cluster if their molecular similarity was greater than 60%, as determined using the Dice coefficient (Dc)(83).

### Testing iron-binding activity by CAS test experiment

We initially screened 48 small molecules from the commercial natural product library for potential siderophore iron-chelating activity, alongside 8 small molecules that lacked such activity, using our developed method. Subsequently, we procured these small molecules individually (Table S6) The small molecules were then prepared by dissolving them in sterilized deionized water to achieve a final concentration of 1 mg/ml in an aqueous solution.

These aqueous solutions were employed to assess the iron-binding activity through the CAS test experiment(59). Briefly, the liquid form of the CAS assay was employed, wherein 100 µl of the small molecule aqueous solution (four biological replicates for all small molecules) or sterilized deionized water as a control reference was added to 100 µl of the CAS assay solution in a 96-well plate. Following a static incubation of 2 hours at room temperature, the OD630 readings of the small molecule aqueous solution and sterilized deionized water and dimethyl sulfoxide (DMSO) controls were measured using a SpectraMax M5 plate reader at room temperature. Small molecules exhibiting siderophore iron-chelating activity would induce a color change in the CAS medium, leading to decreased OD630 measurements. Consequently, the ability of small molecules to chelate iron can be quantified using the OD630 measurements.

## Acknowledgements

We thank Professor Luhua Lai for her insightful comments and suggestions.

## Data Availability Statements

The data underlying this article are available in Zenodo, at https://zenodo.org/doi/10.5281/zenodo.10369626, and can be accessed with 10.5281/zenodo.10369626. The codes underlying this article are available in GitHub at https://github.com/RuolinHe/SIDERITE. The database is available at http://siderite.bdainformatics.org.

## Reference

1. Puig, S., Ramos-Alonso, L., Romero, A.M. and Martinez-Pastor, M. (2017) The elemental role of iron in DNA synthesis and repair. Metallomics, 9, 1483–1500.

2. Read, A.D., Bentley, R.E.T., Archer, S.L. and Dunham-Snary, K.J. (2021) Mitochondrial iron-sulfur clusters: Structure, function, and an emerging role in vascular biology. Redox Biol, 47.

3. Hider, R.C. and Kong, X.L. (2010) Chemistry and biology of siderophores. Nat Prod Rep, 27, 637–657.

4. He, R., Zhang, J., Shao, Y., Gu, S., Song, C., Qian, L., Yin, W.-B. and Li, Z. (2023) Knowledge-guided data mining on the standardized architecture of NRPS: Subtypes, novel motifs, and sequence entanglements. PLOS Computational Biology, 19, e1011100.

5. Kramer, J., Özkaya, Ö. and Kümmerli, R. (2020) Bacterial siderophores in community and host interactions. Nature Reviews Microbiology, 18, 152–163.

6. Page, M.G.P. (2019) The Role of Iron and Siderophores in Infection, and the Development of Siderophore Antibiotics. Clinical Infectious Diseases, 69, S529–S537.

7. Ahmed, E. and Holmström, S.J.M. (2014) Siderophores in environmental research: roles and applications. Microbial Biotechnology, 7, 196–208.

8. Barry, S.M. and Challis, G.L. (2009) Recent advances in siderophore biosynthesis. Current Opinion in Chemical Biology, 13, 205–215.

9. Stow, P.R., Reitz, Z.L., Johnstone, T.C. and Butler, A. (2021) Genomics-driven discovery of chiral triscatechol siderophores with enantiomeric Fe(iii) coordination. Chemical Science, 12, 12485–12493.

10. Hai, Y., Jenner, M. and Tang, Y. (2020) Fungal siderophore biosynthesis catalysed by an iterative nonribosomal peptide synthetase. Chemical Science, 11, 11525–11530.

11. Carroll, C.S. and Moore, M.M. (2018) Ironing out siderophore biosynthesis: a review of non-ribosomal peptide synthetase (NRPS)-independent siderophore synthetases. Crit Rev Biochem Mol, 53, 356–381.

12. Gauglitz, J.M. and Butler, A. (2013) Amino acid variability in the peptide composition of a suite of amphiphilic peptide siderophores from an open ocean Vibrio species. JBIC Journal of Biological Inorganic Chemistry, 18, 489–497.

13. Gao, Y., Walt, C., Bader, C.D. and Müller, R. (2023) Genome-Guided Discovery of the Myxobacterial Thiolactone-Containing Sorangibactins. ACS Chemical Biology, 18, 924–932.

14. Grosse, C., Brandt, N., Van Antwerpen, P., Wintjens, R. and Matthijs, S. (2023) Two new siderophores produced by Pseudomonas sp. NCIMB 10586: The anti-oomycete non-ribosomal peptide synthetase-dependent mupirochelin and the NRPS-independent triabactin. Frontiers in Microbiology, 14.

15. Rehm, K., Vollenweider, V., Gu, S., Friman, V.-P., Kümmerli, R., Wei, Z. and Bigler, L. (2023) Chryseochelins—structural characterization of novel citrate-based siderophores produced by plant protecting Chryseobacterium spp. Metallomics, 15.

16. Hermenau, R., Ishida, K., Gama, S., Hoffmann, B., Pfeifer-Leeg, M., Plass, W., Mohr, J.F., Wichard, T., Saluz, H.-P. and Hertweck, C. (2018) Gramibactin is a bacterial siderophore with a diazeniumdiolate ligand system. Nature Chemical Biology, 14, 841–843.

17. Hermenau, R., Mehl, J.L., Ishida, K., Dose, B., Pidot, S.J., Stinear, T.P. and Hertweck, C. (2019) Genomics-Driven Discovery of NO-Donating Diazeniumdiolate Siderophores in Diverse Plant-Associated Bacteria. Angewandte Chemie International Edition, 58, 13024–13029.

18. Martinet, L., Naômé, A., Deflandre, B., Maciejewska, M., Tellatin, D., Tenconi, E., Smargiasso, N., Pauw, E.d., Wezel, G.P.v. and Rigali, S. (2019) A Single Biosynthetic Gene Cluster Is Responsible for the Production of Bagremycin Antibiotics and Ferroverdin Iron Chelators. mBio, 10, e01230–01219.

19. Martinet, L., Baiwir, D., Mazzucchelli, G. and Rigali, S. (2022) Structure of New Ferroverdins Recruiting Unconventional Ferrous Iron Chelating Agents. Biomolecules, 12, 752.

20. Sorokina, M., Merseburger, P., Rajan, K., Yirik, M.A. and Steinbeck, C. (2021) COCONUT online: Collection of Open Natural Products database. Journal of Cheminformatics, 13, 2.

21. Rutz, A., Sorokina, M., Galgonek, J., Mietchen, D., Willighagen, E., Gaudry, A., Graham, J.G., Stephan, R., Page, R., Vondrášek, J. et al. (2022) The LOTUS initiative for open knowledge management in natural products research. eLife, 11, e70780.

22. Gallo, K., Kemmler, E., Goede, A., Becker, F., Dunkel, M., Preissner, R. and Banerjee, P. (2022) SuperNatural 3.0—a database of natural products and natural product-based derivatives. Nucleic Acids Research, 51, D654–D659.

23. Weininger, D. (1988) SMILES, a chemical language and information system. 1. Introduction to methodology and encoding rules. Journal of Chemical Information and Computer Sciences, 28, 31–36.

24. Hirohara, M., Saito, Y., Koda, Y., Sato, K. and Sakakibara, Y. (2018) Convolutional neural network based on SMILES representation of compounds for detecting chemical motif. BMC Bioinformatics, 19, 526.

25. Rajan, K., Brinkhaus, H.O., Zielesny, A. and Steinbeck, C. (2020) A review of optical chemical structure recognition tools. Journal of Cheminformatics, 12, 60.

26. Clevert, D.-A., Le, T., Winter, R. and Montanari, F. (2021) Img2Mol – accurate SMILES recognition from molecular graphical depictions. Chemical Science, 12, 14174–14181.

27. Khokhlov, I., Krasnov, L., Fedorov, M.V. and Sosnin, S. (2022) Image2SMILES: Transformer-Based Molecular Optical Recognition Engine**. Chemistry–Methods, 2, e202100069.

28. Figueiredo, A.R.T., Ozkaya, O., Kummerli, R. and Kramer, J. (2022) Siderophores drive invasion dynamics in bacterial communities through their dual role as public good versus public bad. Ecol Lett, 25, 138–150.

29. Poggiali, E., Cassinerio, E., Zanaboni, L. and Cappellini, M.D. (2012) An update on iron chelation therapy. Blood Transfus, 10, 411–422.

30. Gu, S.H., Wei, Z., Shao, Z.Y., Friman, V.P., Cao, K.H., Yang, T.J., Kramer, J., Wang, X.F., Li, M., Mei, X.L. et al. (2020) Competition for iron drives phytopathogen control by natural rhizosphere microbiomes. Nat Microbiol, 5, 1002-+.

31. Gu, S., Yang, T., Shao, Z., Wang, T., Cao, K., Jousset, A., Friman, V.P., Mallon, C., Mei, X., Wei, Z., et al. (2020) Siderophore-Mediated Interactions Determine the Disease Suppressiveness of Microbial Consortia. mSystems, 5.

32. Papanikolaou, G. and Pantopoulos, K. (2005) Iron metabolism and toxicity. Toxicology and Applied Pharmacology, 202, 199–211.

33. Hughes, C.E., Coody, T.K., Jeong, M.Y., Berg, J.A., Winge, D.R. and Hughes, A.L. (2020) Cysteine Toxicity Drives Age-Related Mitochondrial Decline by Altering Iron Homeostasis. Cell, 180, 296-+.

34. Sun, H.Y., Zhou, Y., Skaro, M.F., Wu, Y.R., Qu, Z.X., Mao, F.L., Zhao, S.W. and Xu, Y. (2020) Metabolic Reprogramming in Cancer Is Induced to Increase Proton Production. Cancer Res, 80, 1143–1155.

35. Roemhild, K., von Maltzahn, F., Weiskirchen, R., Knuchel, R., von Stillfried, S. and Lammers, T. (2021) Iron metabolism: pathophysiology and pharmacology. Trends Pharmacol Sci, 42, 640–656.

36. Reitz, Z.L., Hardy, C.D., Suk, J., Bouvet, J. and Butler, A. (2019) Genomic analysis of siderophore β-hydroxylases reveals divergent stereocontrol and expands the condensation domain family. Proceedings of the National Academy of Sciences, 116, 19805–19814.

37. Ferreira, D., Seca, A.M.L., Pinto, D.C.G.A. and Silva, A.M.S. (2016) Targeting human pathogenic bacteria by siderophores: A proteomics review. J Proteomics, 145, 153–166.

38. Ye, L.M., Ballet, S., Hildebrand, F., Laus, G., Guillemyn, K., Raes, J., Matthijs, S., Martins, J. and Cornelis, P. (2013) A combinatorial approach to the structure elucidation of a pyoverdine siderophore produced by a Pseudomonas putida isolate and the use of pyoverdine as a taxonomic marker for typing P-putida subspecies. Biometals, 26, 561–575.

39. Zhao, H., Yang, Y., Wang, S.Q., Yang, X., Zhou, K.C., Xu, C.L., Zhang, X.Y., Fan, J.J., Hou, D.Y., Li, X.X. et al. (2022) NPASS database update 2023: quantitative natural product activity and species source database for biomedical research. Nucleic Acids Research.

40. van Santen, J.A., Poynton, E.F., Iskakova, D., McMann, E., Alsup, T.A., Clark, T.N., Fergusson, C.H., Fewer, D.P., Hughes, A.H., McCadden, C.A., et al. (2022) The Natural Products Atlas 2.0: a database of microbially-derived natural products. Nucleic Acids Research, 50, D1317–D1323.

41. D’Onofrio, A., Crawford, J.M., Stewart, E.J., Witt, K., Gavrish, E., Epstein, S., Clardy, J. and Lewis, K. (2010) Siderophores from Neighboring Organisms Promote the Growth of Uncultured Bacteria. Chem Biol, 17, 254–264.

42. Maglangit, F., Tong, M.H., Jaspars, M., Kyeremeh, K. and Deng, H. (2019) Legonoxamines A-B, two new hydroxamate siderophores from the soil bacterium, Streptomyces sp. MA37. Tetrahedron Lett, 60, 75–79.

43. Wilson, M.K., Abergel, R.J., Raymond, K.N., Arceneaux, J.E.L. and Byers, B.R. (2006) Siderophores of Bacillus anthracis, Bacillus cereus, and Bacillus thuringiensis. Biochem Bioph Res Co, 348, 320–325.

44. Zajdowicz, S., Haller, J.C., Krafft, A.E., Hunsucker, S.W., Mant, C.T., Duncan, M.W., Hodges, R.S., Jones, D.N.M. and Holmes, R.K. (2012) Purification and Structural Characterization of Siderophore (Corynebactin) from Corynebacterium diphtheriae. Plos One, 7.

45. Patzer, S.I. and Braun, V. (2010) Gene Cluster Involved in the Biosynthesis of Griseobactin, a Catechol-Peptide Siderophore of Streptomyces sp ATCC 700974. J Bacteriol, 192, 426–435.

46. Matsuo, Y., Kanoh, K., Jang, J.H., Adachi, K., Matsuda, S., Miki, O., Kato, T. and Shizuri, Y. (2011) Streptobactin, a Tricatechol-Type Siderophore from Marine-Derived Streptomyces sp YM5-799. J Nat Prod, 74, 2371–2376.

47. Carrero, M.I.G., Sangari, F.J., Aguero, J. and Lobo, J.M.G. (2002) Brucella abortus strain 2308 produces brucebactin, a highly efficient catecholic siderophore. Microbiol-Sgm, 148, 353–360.

48. Miller, A.L., Li, S.R., Eichhorn, C.D., Zheng, Y.B. and Du, L.C. (2023) Identification and Biosynthetic Study of the Siderophore Lysochelin in the Biocontrol Agent Lysobacter enzymogenes. J Agr Food Chem.

49. Raines, D.J., Moroz, O.V., Blagova, E.V., Turkenburg, J.P., Wilson, K.S. and Duhme-Klair, A.-K. (2016) Bacteria in an intense competition for iron: Key component of the *Campylobacter jejuni* iron uptake system scavenges enterobactin hydrolysis product. Proceedings of the National Academy of Sciences, 113, 5850–5855.

50. Kummerli, R., Schiessl, K.T., Waldvogel, T., McNeill, K. and Ackermann, M. (2014) Habitat structure and the evolution of diffusible siderophores in bacteria. Ecol Lett, 17, 1536–1544.

51. Francoeur, P.G. and Koes, D.R. (2021) SolTranNet-A Machine Learning Tool for Fast Aqueous Solubility Prediction. J Chem Inf Model, 61, 2530–2536.

52. Evans, R., Dal Poggetto, G., Nilsson, M. and Morris, G.A. (2018) Improving the Interpretation of Small Molecule Diffusion Coefficients. Anal Chem, 90, 3987–3994.

53. Völker, C. and Wolf-Gladrow, D.A. (1999) Physical limits on iron uptake mediated by siderophores or surface reductases. Marine Chemistry, 65, 227–244.

54. Demain, A.L. (1992) Microbial Secondary Metabolism - a New Theoretical Frontier for Academia, a New Opportunity for Industry. Ciba F Symp, 171, 3–23.

55. Sinsabaugh, R.L., Hill, B.H. and Shah, J.J.F. (2009) Ecoenzymatic stoichiometry of microbial organic nutrient acquisition in soil and sediment. Nature, 462, 795–U117.

56. Chen, Y.L., Chen, L.Y., Peng, Y.F., Ding, J.Z., Li, F., Yang, G.B., Kou, D., Liu, L., Fang, K., Zhang, B.B. et al. (2016) Linking microbial C:N:P stoichiometry to microbial community and abiotic factors along a 3500-km grassland transect on the Tibetan Plateau. Global Ecol Biogeogr, 25, 1416–1427.

57. Probst, D. and Reymond, J.L. (2020) Visualization of very large high-dimensional data sets as minimum spanning trees. Journal of Cheminformatics, 12.

58. Vivanco, J.M., Bais, H.P., Stermitz, F.R., Thelen, G.C. and Callaway, R.M. (2004) Biogeographical variation in community response to root allelochemistry: novel weapons and exotic invasion. Ecol Lett, 7, 285–292.

59. Schwyn, B. and Neilands, J.B. (1987) Universal chemical assay for the detection and determination of siderophores. Anal Biochem, 160, 47–56.

60. Fazary, A.E., Taha, M. and Ju, Y.H. (2009) Iron Complexation Studies of Gallic Acid. J Chem Eng Data, 54, 35–42.

61. Matthijs, S., Baysse, C., Koedam, N., Tehrani, K.A., Verheyden, L., Budzikiewicz, H., Schafer, M., Hoorelbeke, B., Meyer, J.M., De Greve, H., et al. (2004) The Pseudomonas siderophore quinolobactin is synthesized from xanthurenic acid, an intermediate of the kynurenine pathway. Molecular Microbiology, 52, 371–384.

62. Barraza, M.L., Coppock, C.E., Brooks, K.N., Wilks, D.L., Saunders, R.G. and Latimer, G.W. (1991) Iron Sulfate and Feed Pelleting to Detoxify Free Gossypol in Cottonseed Diets for Dairy-Cattle. J Dairy Sci, 74, 3457–3467.

63. Mossialos, D., Meyer, J.-M., Budzikiewicz, H., Wolff, U., Koedam, N., Baysse, C., Anjaiah, V. and Cornelis, P. (2000) Quinolobactin, a New Siderophore of *Pseudomonas fluorescens* ATCC 17400, the Production of Which Is Repressed by the Cognate Pyoverdine. Applied and Environmental Microbiology, 66, 487–492.

64. Matthijs, S., Tehrani, K.A., Laus, G., Jackson, R.W., Cooper, R.M. and Cornelis, P. (2007) Thioquinolobactin, a Pseudomonas siderophore with antifungal and anti-Pythium activity. Environ Microbiol, 9, 425–434.

65. 石井, 永., 陳, 益., 赤池, 美., 石川, 勉. and 盧, 盛. (1982) ミカン科植物成分の研究(第 44 報)台湾産ネワタノキ Xanthoxylum integrifoliolum(MERR.)MERR.(Fagara integrifoliola MERR.)の成分検索 その 1 根木質部の成分. 藥學雜誌, 102, 182-195.

66. Chen, J.-J., Wang, T.-Y. and Hwang, T.-L. (2008) Neolignans, a Coumarinolignan, Lignan Derivatives, and a Chromene: Anti-inflammatory Constituents from Zanthoxylum avicennae. J Nat Prod, 71, 212–217.

67. Moon, S.-S., Cho, N., Shin, J., Seo, Y., Lee, C.O. and Choi, S.U. (1996) Jineol, a Cytotoxic Alkaloid from the Centipede Scolopendra subspinipes. J Nat Prod, 59, 777–779.

68. Cordero, O.X., Ventouras, L.A., DeLong, E.F. and Polz, M.F. (2012) Public good dynamics drive evolution of iron acquisition strategies in natural bacterioplankton populations. P Natl Acad Sci USA, 109, 20059–20064.

69. Butaite, E., Baumgartner, M., Wyder, S. and Kummerli, R. (2017) Siderophore cheating and cheating resistance shape competition for iron in soil and freshwater Pseudomonas communities. Nature Communications, 8.

70. Ozkaya, O., Balbontin, R., Gordo, I. and Xavier, K.B. (2018) Cheating on Cheaters Stabilizes Cooperation in Pseudomonas aeruginosa. Curr Biol, 28, 2070-+.

71. Shepherdson, E.M.F. and Elliot, M.A. (2022) Cryptic specialized metabolites drive *Streptomyces* exploration and provide a competitive advantage during growth with other microbes. Proceedings of the National Academy of Sciences, 119, e2211052119.

72. Zheng, S.J., Zeng, T., Li, C.T., Chen, B.H., Coley, C.W., Yang, Y.D. and Wu, R.B. (2022) Deep learning driven biosynthetic pathways navigation for natural products with BioNavi-NP. Nature Communications, 13.

73. Isaka, M., Tanticharoen, M., Kongsaeree, P. and Thebtaranonth, Y. (2001) Structures of cordypyridones A-D, antimalarial N-hydroxy- and N-methoxy-2-pyridones from the insect pathogenic fungus Cordyceps nipponica. J Org Chem, 66, 4803–4808.

74. Pita-Grisanti, V., Chasser, K., Sobol, T. and Cruz-Monserrate, Z. (2022) Understanding the Potential and Risk of Bacterial Siderophores in Cancer. Front Oncol, 12.

75. Ganz, T. and Nemeth, E. (2015) Iron homeostasis in host defence and inflammation. Nat Rev Immunol, 15, 500–510.

76. Nairz, M. and Weiss, G. (2020) Iron in infection and immunity. Mol Aspects Med, 75.

77. Chuljerm, H., Deeudom, M., Fucharoen, S., Mazzacuva, F., Hider, R.C., Srichairatanakool, S. and Cilibrizzi, A. (2020) Characterization of two siderophores produced by Bacillus megaterium: A preliminary investigation into their potential as therapeutic agents. Bba-Gen Subjects, 1864.

78. Achard, M.E.S., Chen, K.W.W., Sweet, M.J., Watts, R.E., Schroder, K., Schembri, M.A. and McEwan, A.G. (2013) An antioxidant role for catecholate siderophores in Salmonella. Biochem J, 454, 543–549.

79. Peralta, D.R., Adler, C., Corbalan, N.S., Garcia, E.C.P., Pomares, M.F. and Vincent, P.A. (2016) Enterobactin as Part of the Oxidative Stress Response Repertoire. Plos One, 11.

80. Hesse, E., O’Brien, S., Tromas, N., Bayer, F., Lujan, A.M., van Veen, E.M., Hodgson, D.J. and Buckling, A. (2018) Ecological selection of siderophore-producing microbial taxa in response to heavy metal contamination. Ecol Lett, 21, 117–127.

81. Roskova, Z., Skarohlid, R. and McGachy, L. (2022) Siderophores: an alternative bioremediation strategy? Sci Total Environ, 819, 153144.

82. Ricart, E., Leclere, V., Flissi, A., Mueller, M., Pupin, M. and Lisacek, F. (2019) rBAN: retro-biosynthetic analysis of nonribosomal peptides. Journal of Cheminformatics, 11.

83. Maggiora, G., Vogt, M., Stumpfe, D. and Bajorath, J. (2014) Molecular similarity in medicinal chemistry. J Med Chem, 57, 3186–3204.

