## Supplementary Figure S1-S18 for "SIDERITE: Unveiling Hidden Siderophore Diversity in the Chemical Space Through Digital Exploration"

#### The legend of Supplementary Materials

Table S1. 872 siderophore information records.
Table S2. Detailed information on 649 siderophores with unique structures.
Table S3. 15 functional group structures used in potential siderophore search.
Table S4. 8 modified siderophore functional groups in the negative control.
Table S5. Detailed information on 48 molecules with potential iron-binding activities in the CAS assay test.
Table S6. The raw OD630 values of 48 molecules with potential iron-binding activities in the CAS assay test.
Table S7. Tanimoto similarity matrix between 649 siderophores in SIDERITE and 3,199 molecules with potential iron-binding activities in COCONUT database.

**Fig S1**

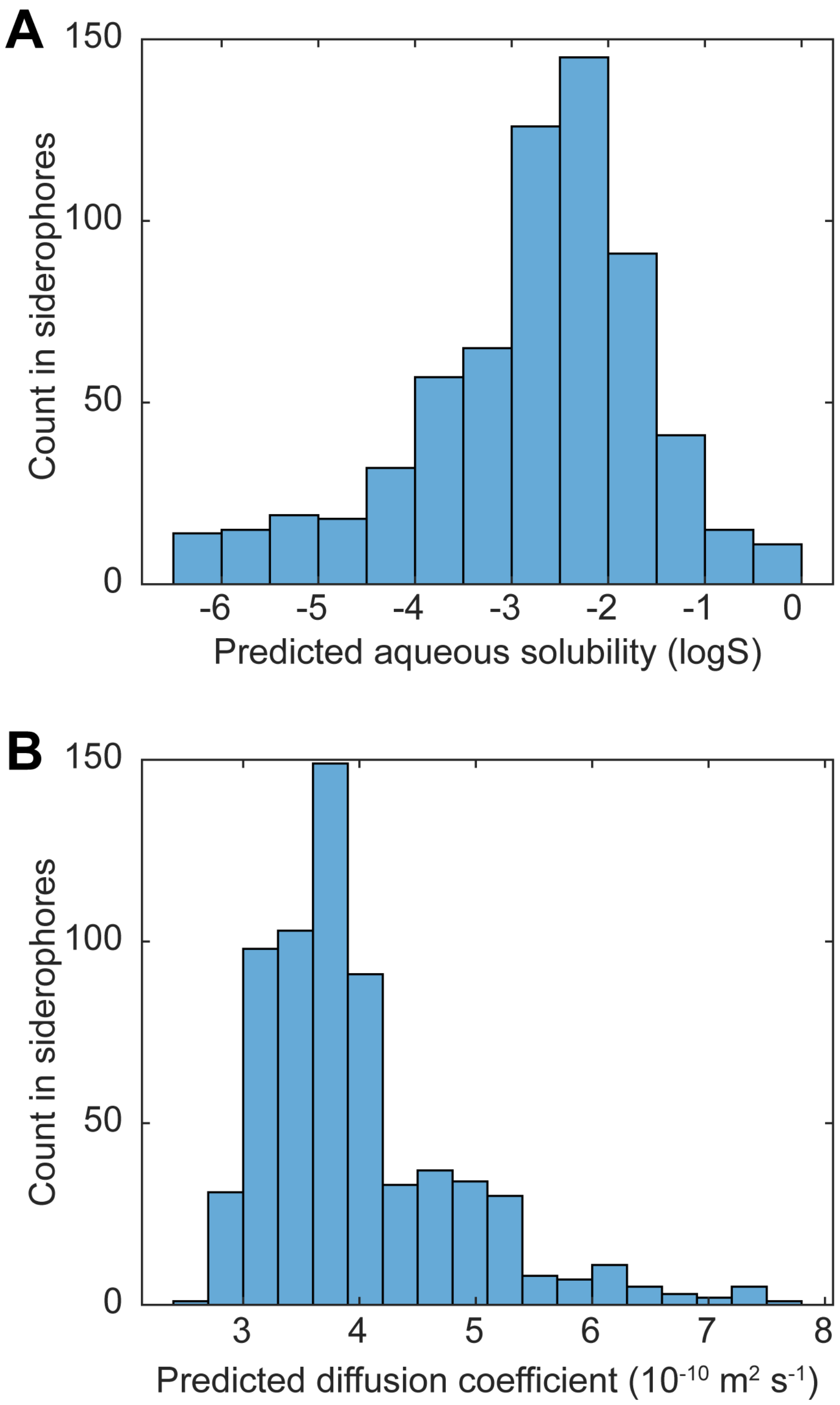


**Fig S1. The predicted properties of 649 siderophores**

A. The predicted aqueous solubility. The unit is log10 of solubility in the water.

B. The predicted diffusion coefficient. The temperature is set to 298.15K and the solvent is water.

### Fig S2

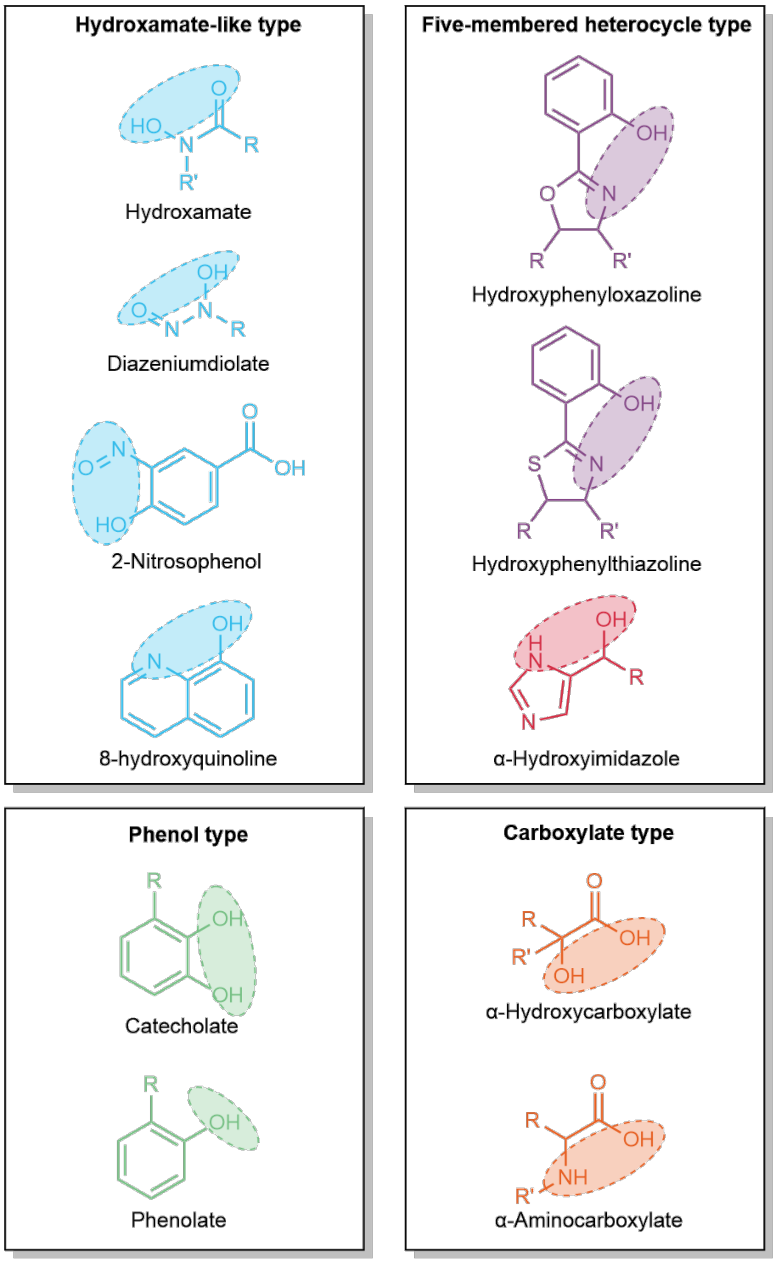


**Fig S2. Known siderophore functional groups (ligands)**

### Fig S3

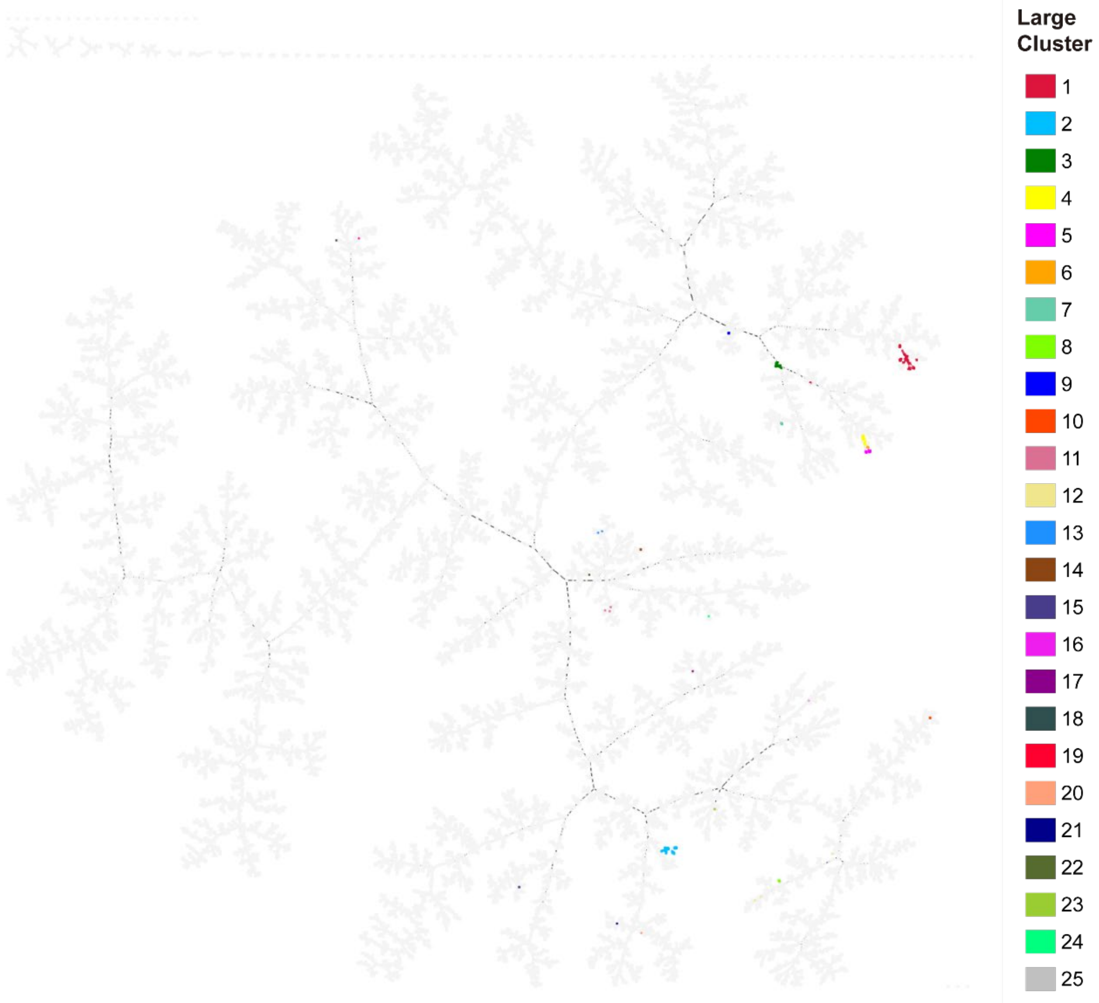


**Fig S3. Displaying 25 clusters of 649 siderophores in the COCONUT database by TMAP**

### Fig S4

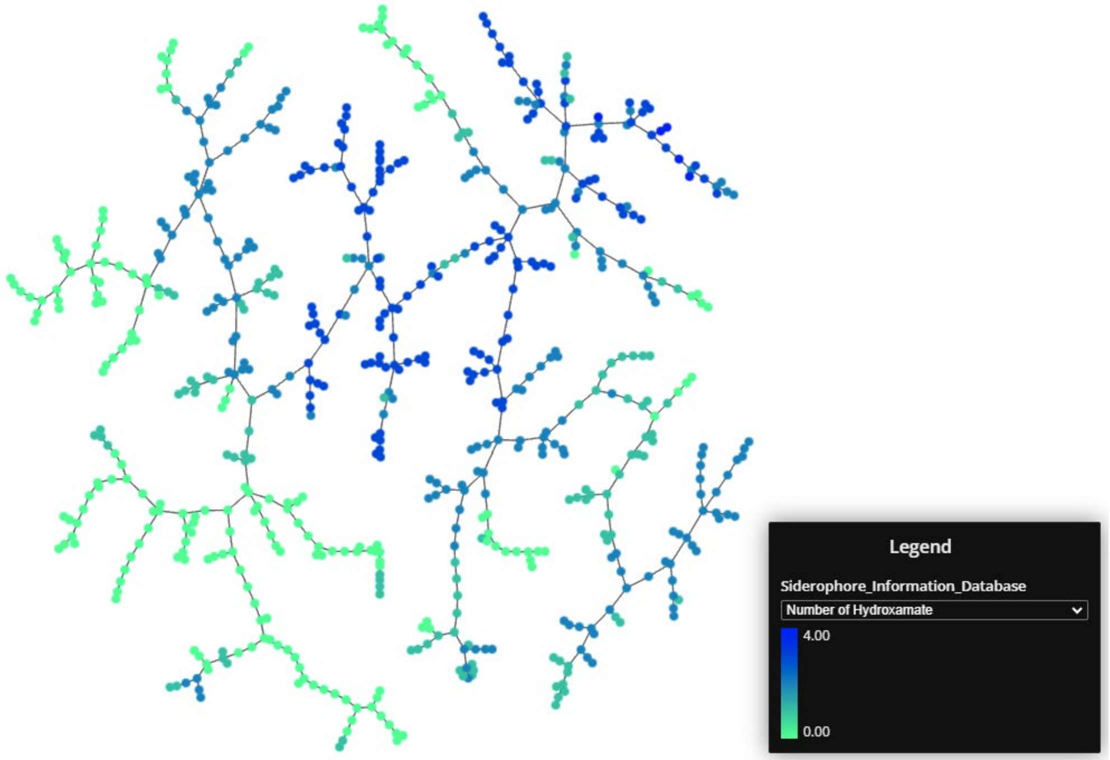


**Fig S4. Visualization of 649 siderophores with functional group hydroxamate number by TAMP**

### Fig S5

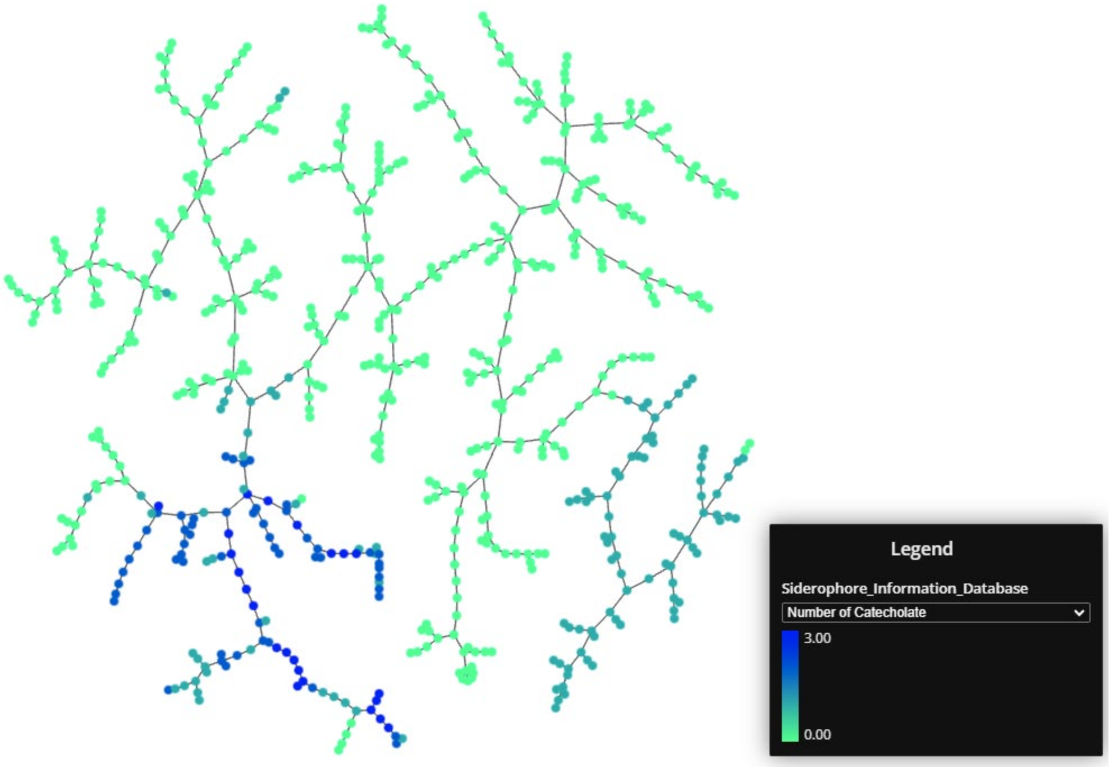


**Fig S5. Visualization of 649 siderophores with functional group catecholate number by TAMP**

#### Fig S6

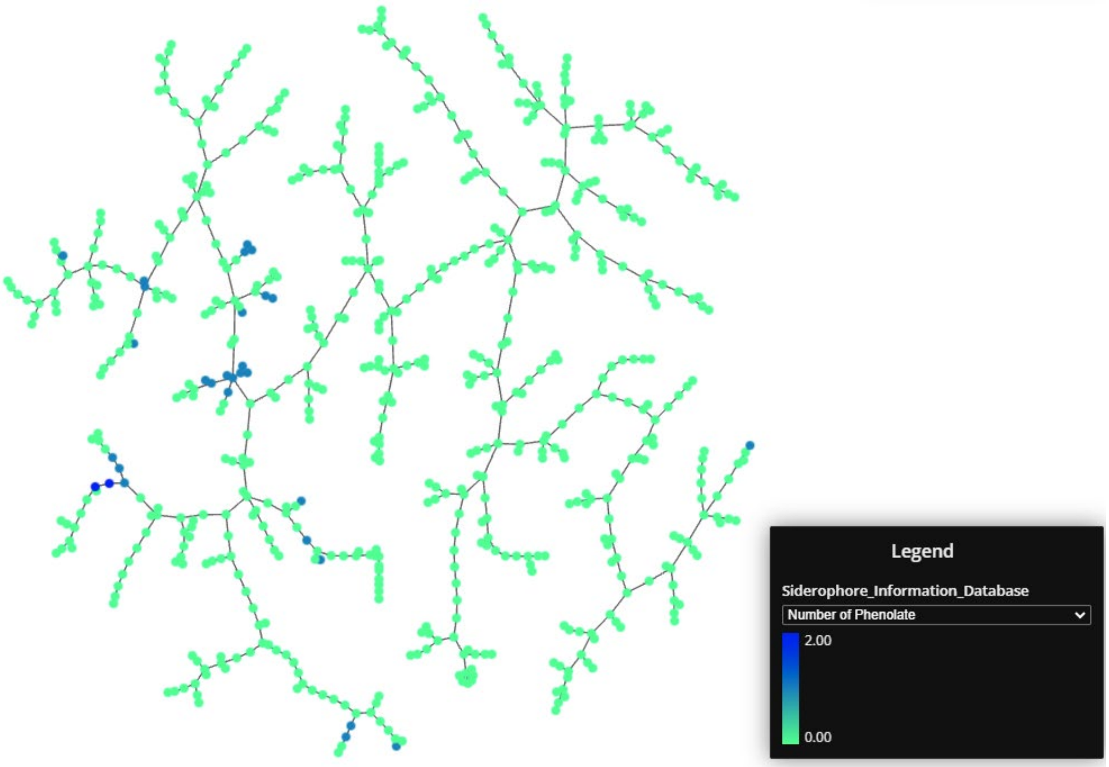


**Fig S6. Visualization of 649 siderophores with functional group phenolate number by TAMP**

### Fig S7

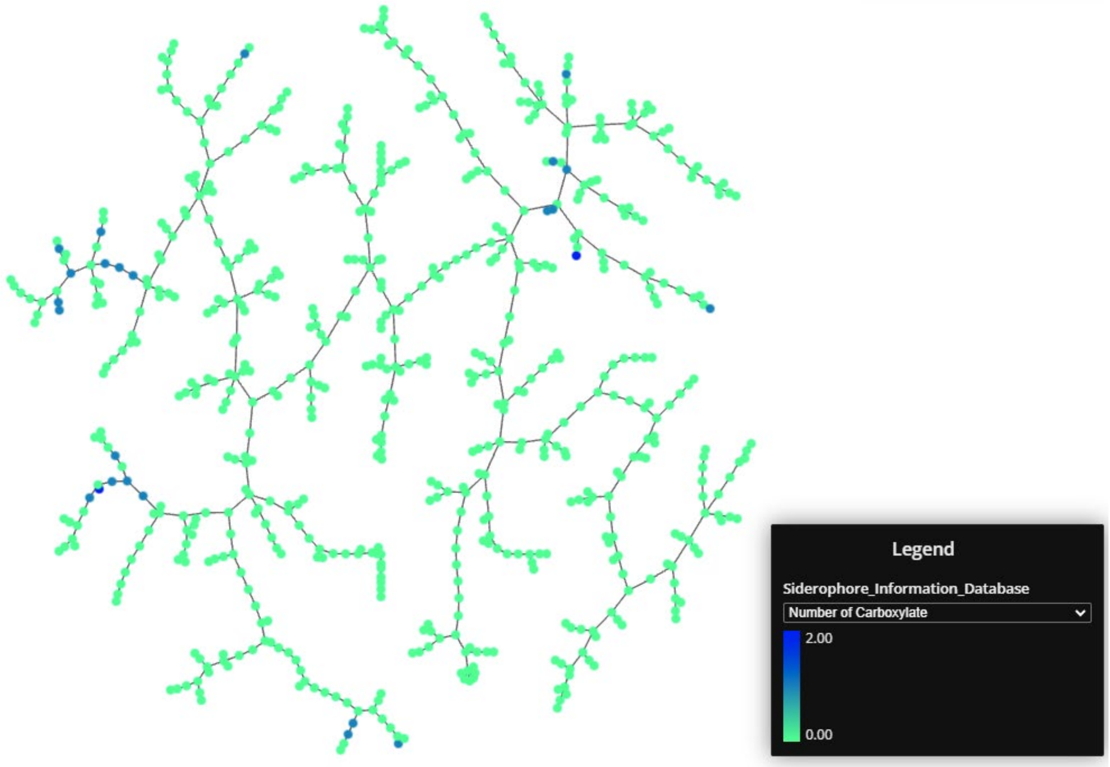


**Fig S7. Visualization of 649 siderophores with functional group carboxylate number by TAMP**

### Fig S8

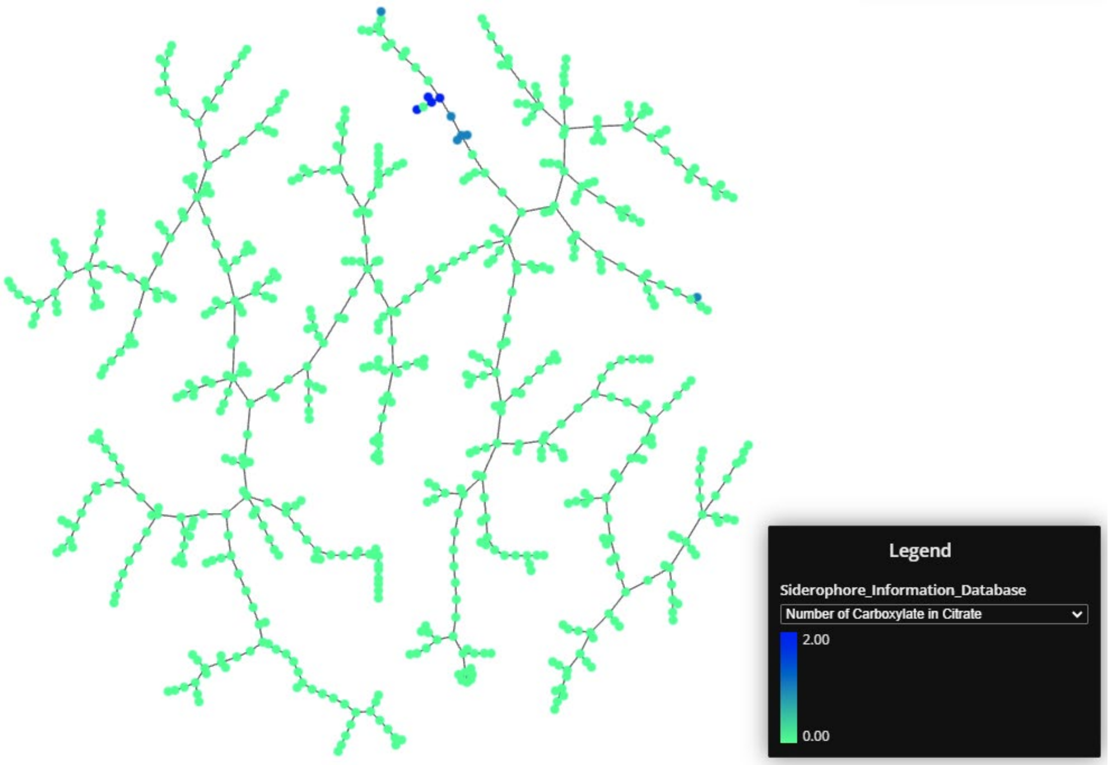


**Fig S8. Visualization of 649 siderophores with functional group carboxylate in citrate number by TAMP**

#### Fig S9

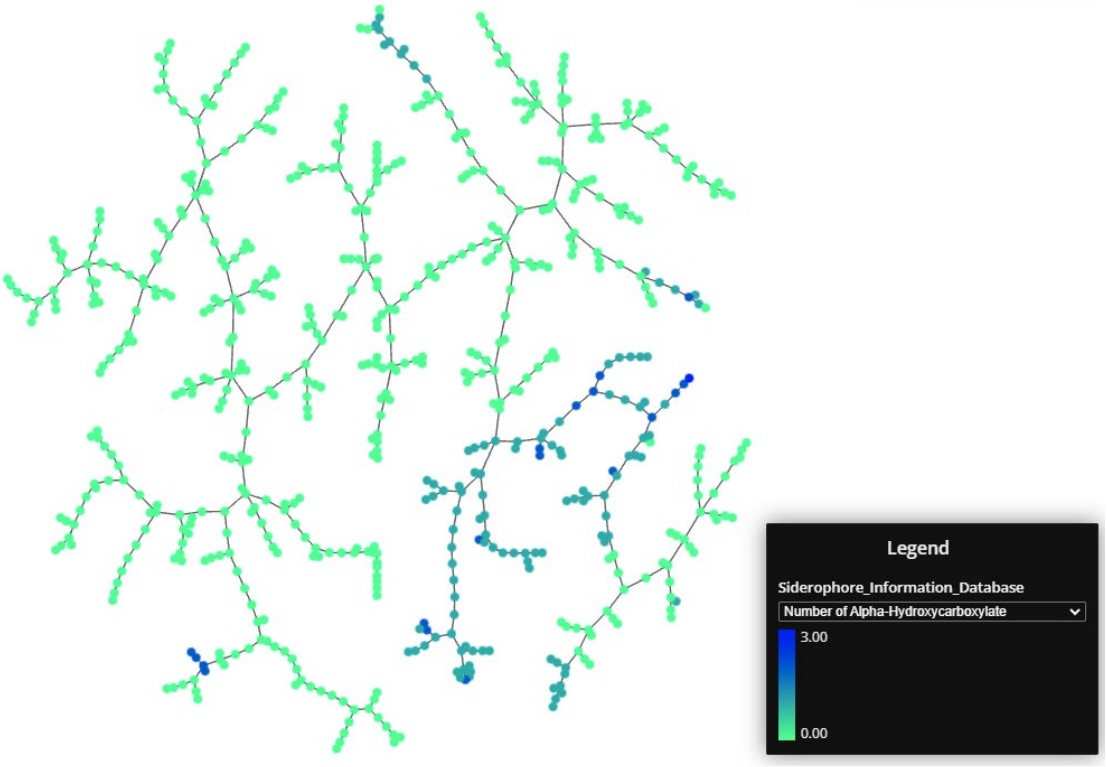


**Fig S9. Visualization of 649 siderophores with functional group alpha-hydroxycarboxylate number by TAMP**

### Fig S10

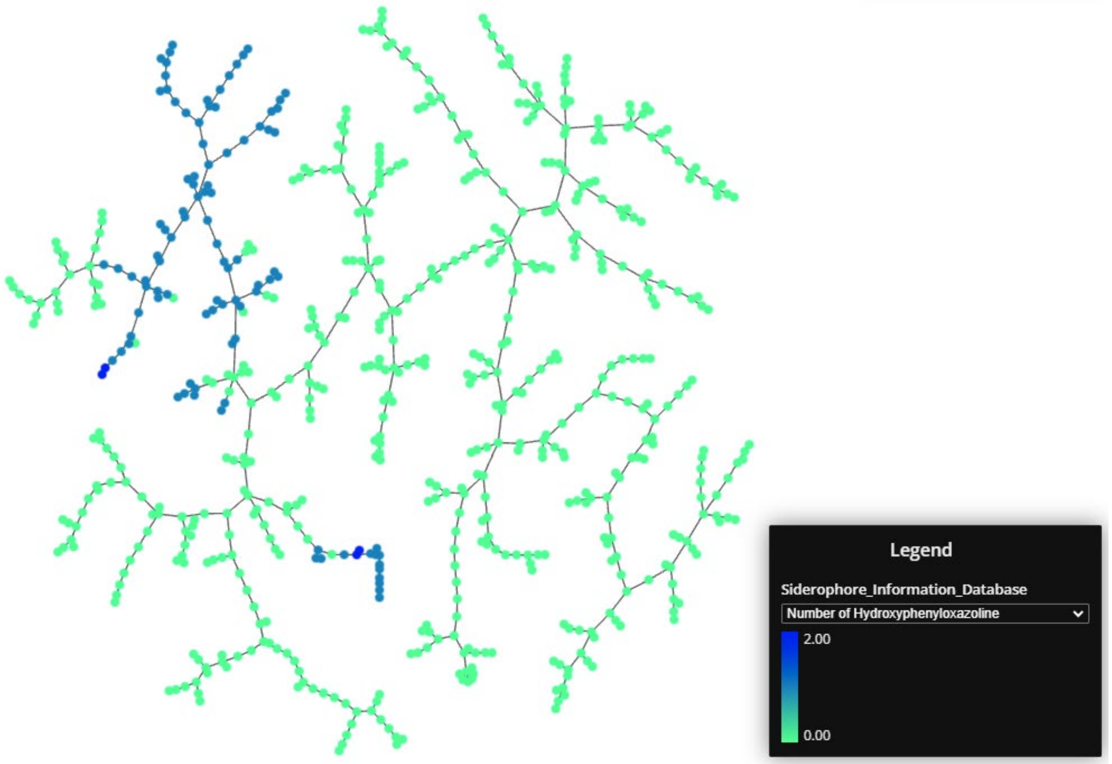


**Fig S10. Visualization of 649 siderophores with functional group hydroxyphenyloxazoline number by TAMP**

### Fig S11

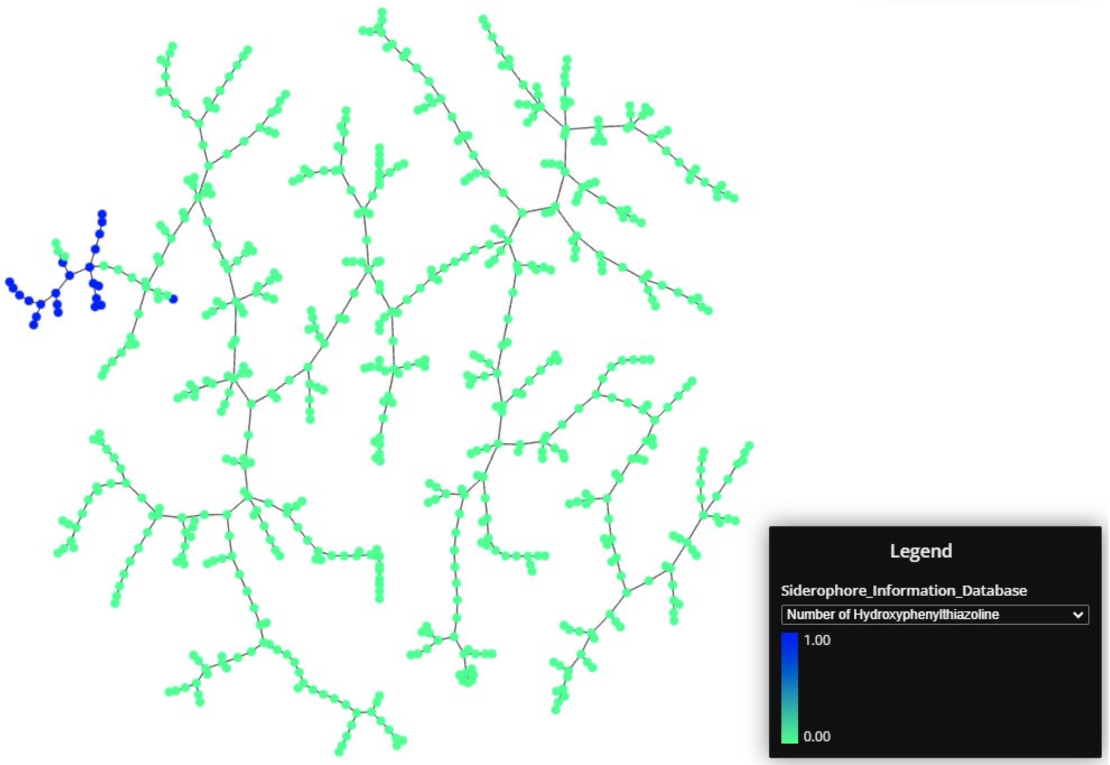


**Fig S11. Visualization of 649 siderophores with functional group hydroxyphenylthiazoline number by TAMP**

### Fig S12

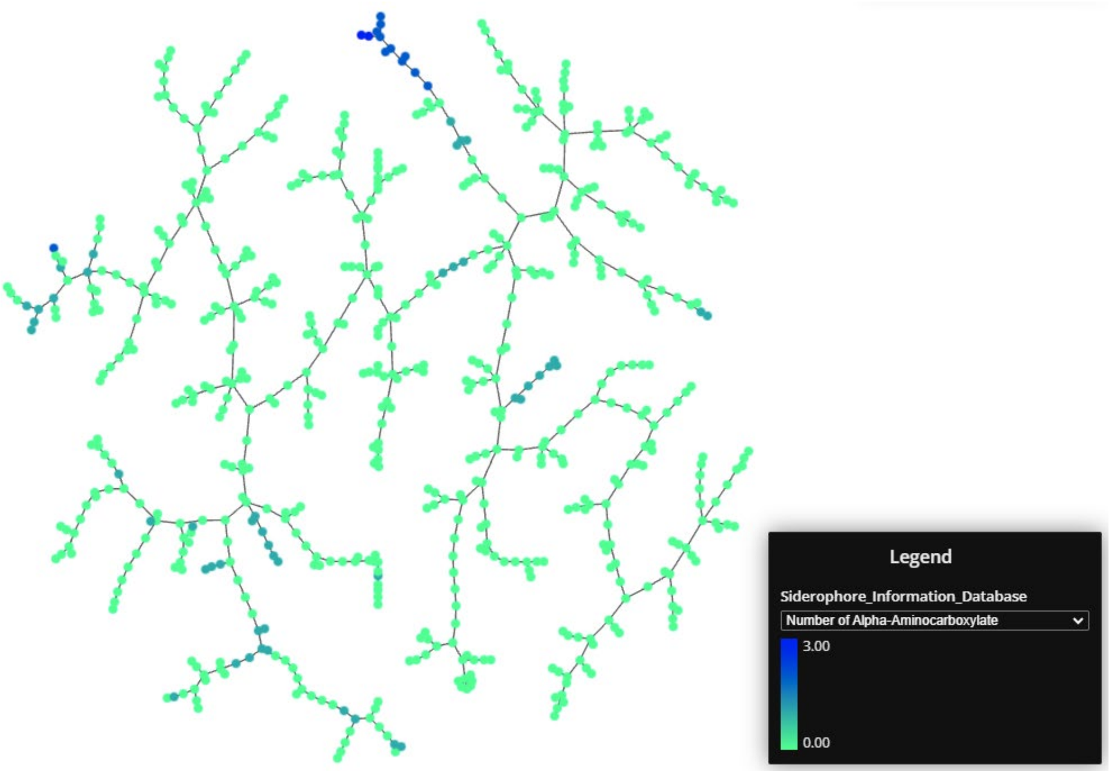


**Fig S12. Visualization of 649 siderophores with functional group alpha-aminocarboxylate number by TAMP**

### Fig S13

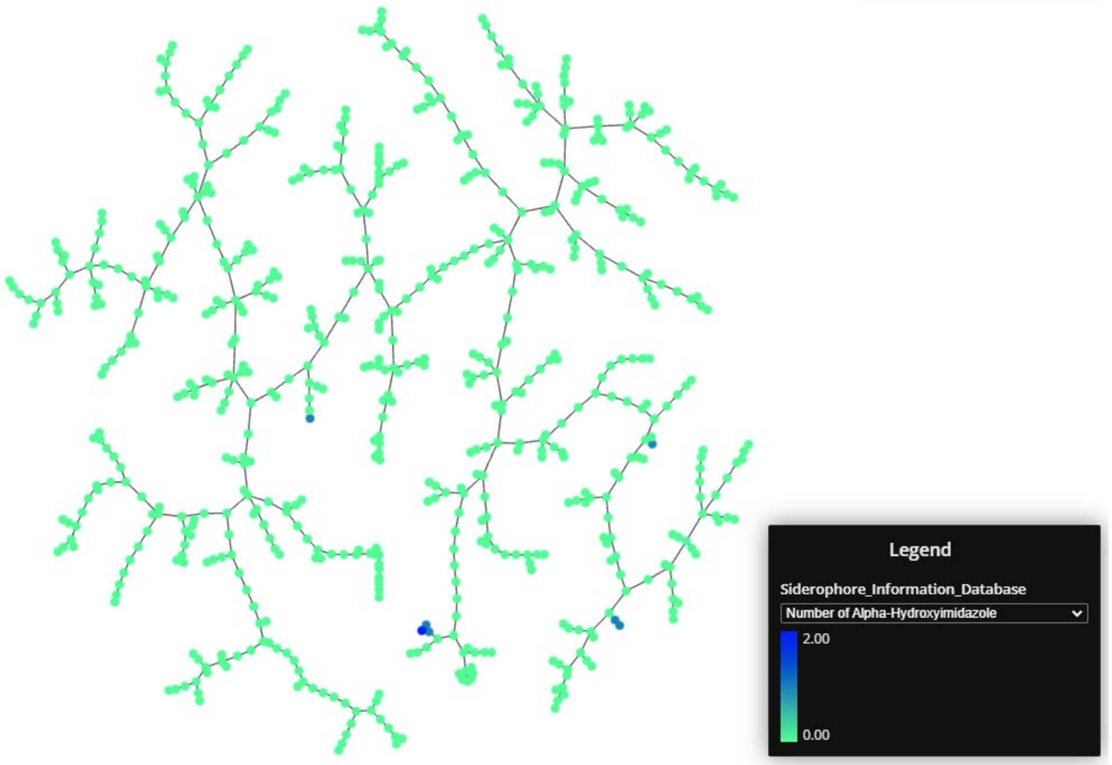


**Fig S13. Visualization of 649 siderophores with functional group alpha-hydroxyimidazole number by TAMP**

#### Fig S14

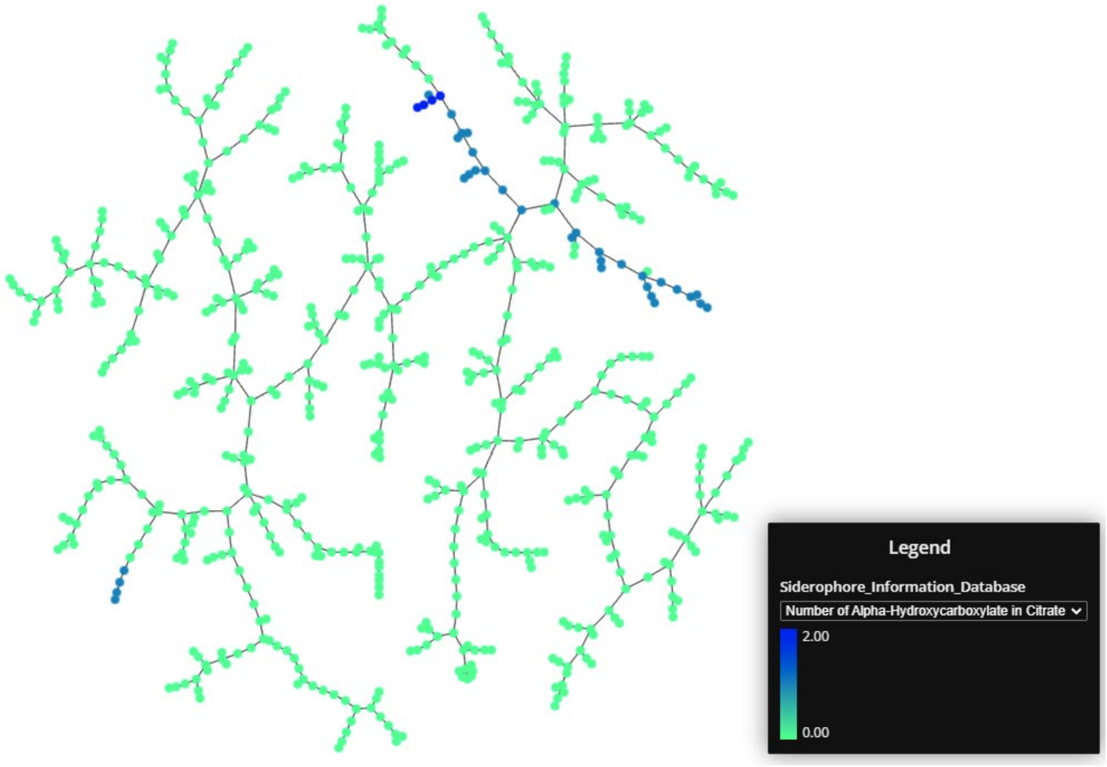


**Fig S14. Visualization of 649 siderophores with functional group alpha-hydroxycarboxylate in citrate number by TAMP**

#### Fig S15

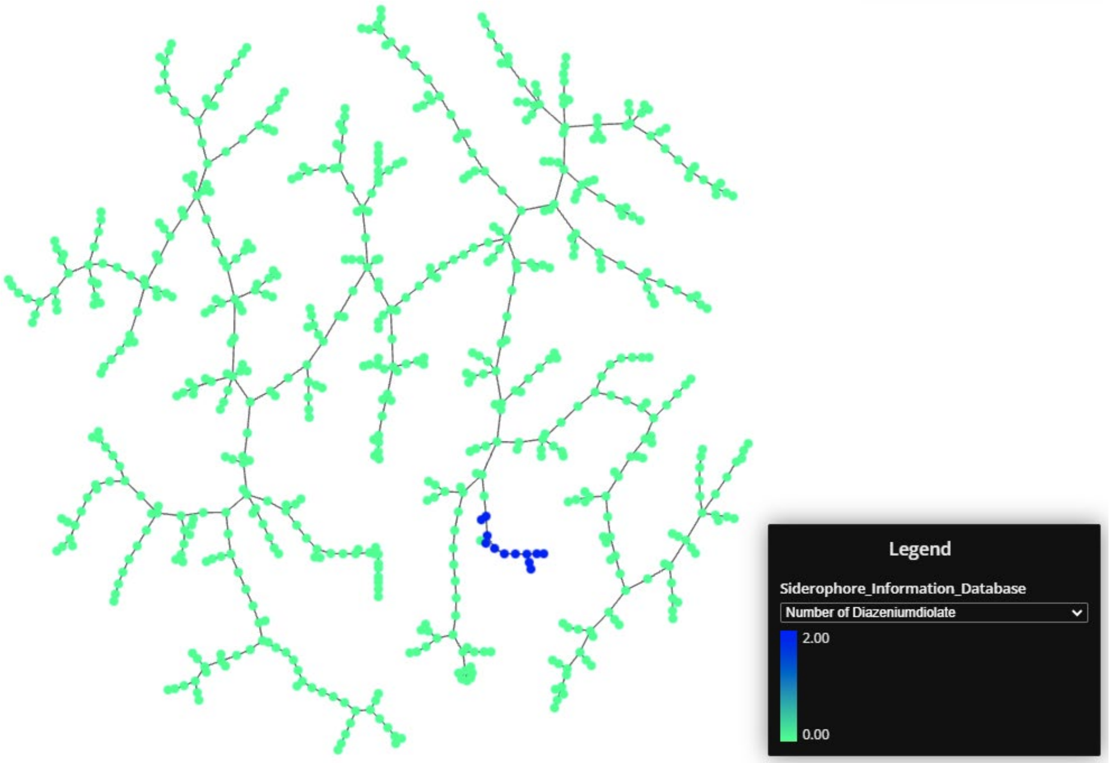


**Fig S15. Visualization of 649 siderophores with functional group diazeniumdiolate number by TAMP**

#### Fig S16

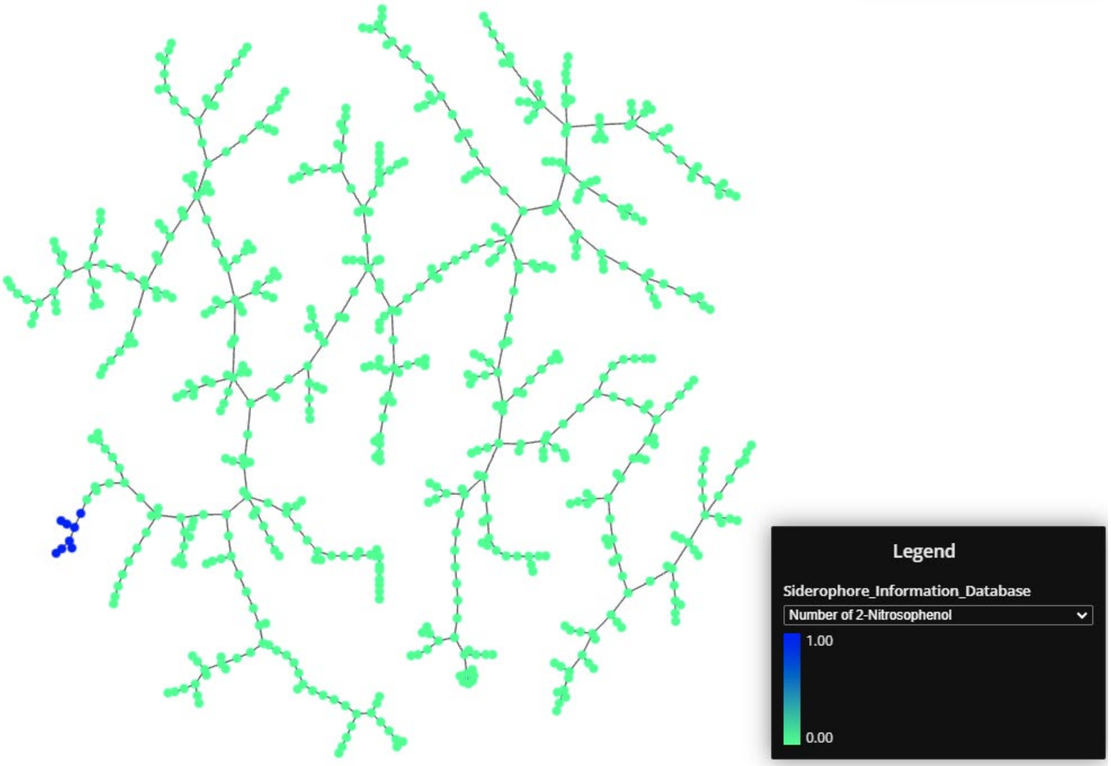


**Fig S16. Visualization of 649 siderophores with functional group 2-nitrosophenol number by TAMP**

### Fig S17

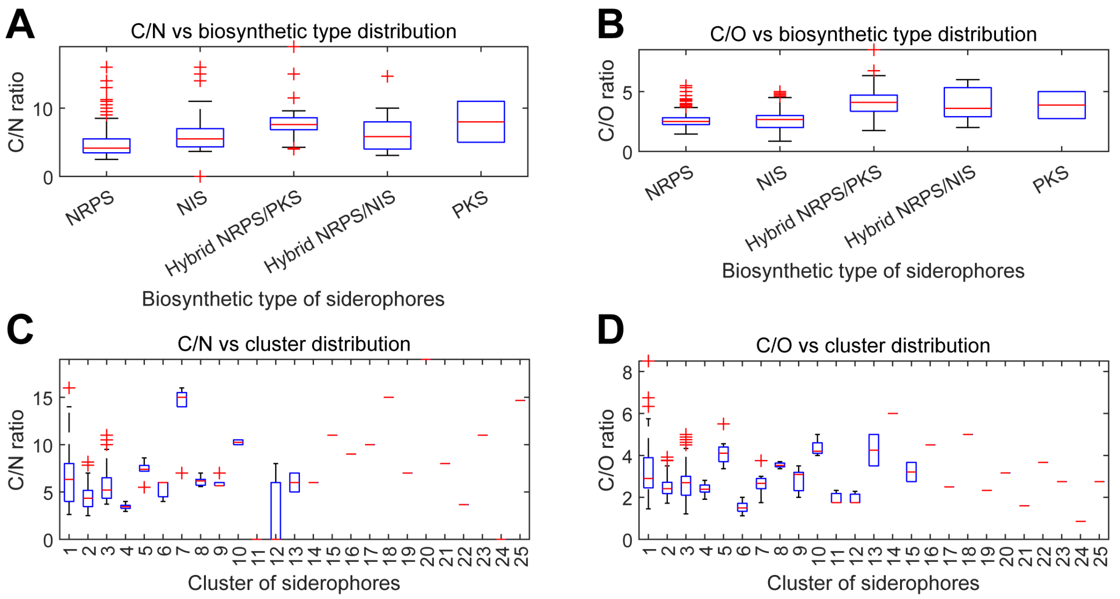


**Fig S17. The distribution of C/N and C/O ratios in the different biosynthetic types and clusters**

**A.** The distribution of C/N ratio between different biosynthetic types of siderophores.

**B.** The distribution of C/O ratio between different biosynthetic types of siderophores.

**C.** The distribution of C/N ratio between different clusters of siderophores.

**D.** The distribution of C/O ratio between different clusters of siderophores.

### Fig S18

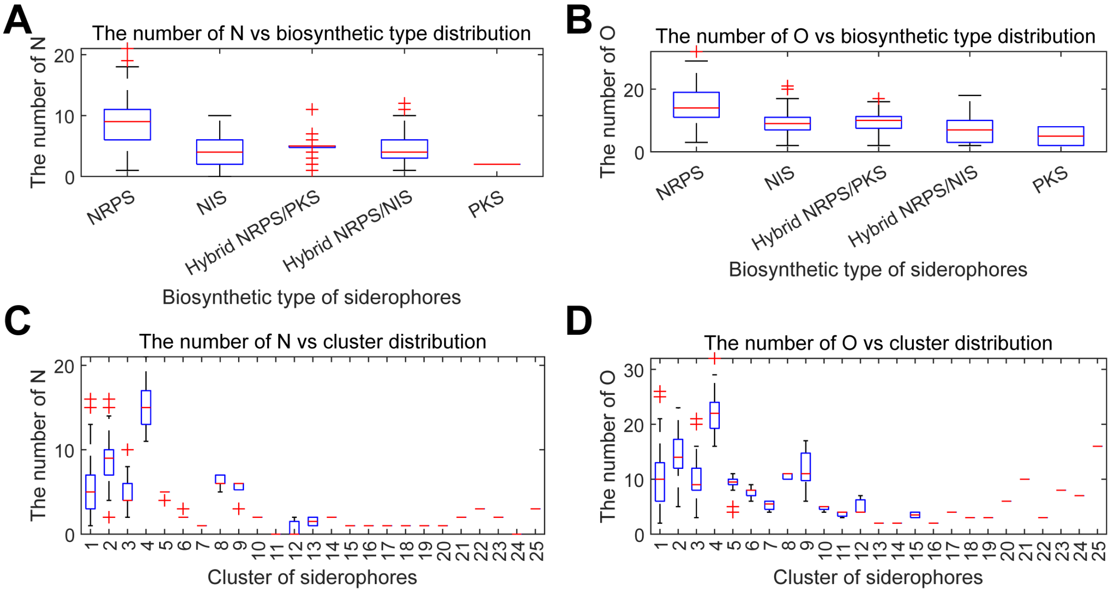


**Fig S18. The distribution of nitrogen atom and oxygen atom numbers in the different biosynthetic types and clusters**

**E.** The distribution of nitrogen atom number between different biosynthetic types of siderophores.

**F.** The distribution of oxygen atom number between different biosynthetic types of siderophores.

**G.** The distribution of nitrogen atom number between different clusters of siderophores.

**H.** The distribution of oxygen atom number between different clusters of siderophores.
